# Europe as a bridgehead in the worldwide invasion history of grapevine downy mildew, *Plasmopara viticola*

**DOI:** 10.1101/2020.09.22.307678

**Authors:** Michael C. Fontaine, Frédéric Labbé, Yann Dussert, Laurent Delière, Sylvie Richart-Cervera, Tatiana Giraud, François Delmotte

## Abstract

Europe is the historical cradle of viticulture, but grapevines have been increasingly threatened by pathogens of American origin. The invasive oomycete *Plasmopara viticola* causes downy mildew, one of the most devastating grapevine diseases worldwide. Despite major economic consequences, its invasion history remains poorly understood. Comprehensive population genetic analyses of ~2000 samples from the most important wine-producing countries revealed very low genetic diversity in invasive downy mildew populations worldwide. All the populations originated from one of five native North American lineages, the one parasitizing wild summer grape. After an initial introduction into Europe, invasive European populations served as a secondary source of introduction into vineyards worldwide, including China, South Africa and, twice independently, Australia. Invasion of Argentina probably represents a tertiary introduction from Australia. Our findings provide a striking example of a global pathogen invasion resulting from secondary dispersal of a successful invasive population. It will help designing quarantine regulations and efficient breeding for resistance against grapevine downy mildew.

## Introduction

Global changes (e.g., climate warming and international exchanges) are favoring increases in the numbers of emerging diseases caused by invasive pathogens on crops worldwide, incurring substantial economic, social and environmental costs (*1*–*3*). Infamous recent examples include the emergence of new races of the stem rust fungus in Eastern Africa (*4*) and of the fungus causing wheat blast disease in Bangladesh (*5*), both threatening wheat production and becoming invasive. Most emerging diseases result from biological invasions bringing the native host-parasite association back together after crop introduction into new areas, or host shifts following pathogen introductions (*6*, *7*). An understanding of the evolutionary processes responsible for emerging crop diseases is important for preventing further devastating biological invasions and for controlling introduced populations. This requires elucidation of the invasion mechanisms, pathways and demographic processes occurring during pathogen invasions (e.g., bottlenecks and hybridization). Important questions include whether pathogen invasions result from host shifts, whether limited genetic variation has been introduced, whether multiple introductions and admixture are required for successful invasions (*7*), and whether the invaded areas are colonized directly from native populations or whether an initial successful invasive population serves as the source for secondary introductions.

Grapevine downy mildew, caused by the oomycete *Plasmopara viticola*, is one of the most damaging diseases of grapevine, and is found in all grape-growing regions of the world (*8*). *Plasmopara viticola* is native to eastern North America (*9*), and was introduced into Europe in the 1870s (*10*), probably with American *Vitis* species used as rootstocks resistant to phylloxera, an insect pest (*Daktulosphaira vitifoliae*) (*9*, *11*). After its first description in France (1878 in Coutras), the disease rapidly reached Southern and Central Europe and was soon reported in nearly all wine-producing countries worldwide (*12*, *13*).

Advances in *P. viticola* -omics (*14*–*19*) and population genetic studies (*20*–*35*) have improved our knowledge of the grapevine downy mildew pathosystem. In its native range, five cryptic species (also called *formae speciales*) have recently been identified in the *P. viticola* species complex, with genetic differentiation and contrasting host ranges on various *Vitis* and related species (*33*, *34*). The five *P. viticola formae speciales* (f. sp.) are found on wild *Vitis* species across North America (*33*, *34*), and it remains unknown which lineages were responsible for grapevine downy mildew invasions in vineyards across the world. In most temperate regions, *P. viticola* populations present widespread footprints of recombination, indicating the occurrence of frequent sexual reproduction (*21*, *24*, *26*, *27*, *29*, *31*, *36*). European invasive populations display little genetic diversity and have a weak but significant population structure at the continental scale (*22*, *25*). Despite some notable advances, however, these studies have been restricted to a small number of countries and were performed with different markers, making it difficult to develop a comprehensive understanding of the pathways of *P. viticola* invasion worldwide.

In this study, we used phylogenetic and population genetic approaches, together with scenario testing by approximate Bayesian computation (ABC), to infer the routes of *P. viticola* invasion worldwide. We analyzed almost 2,000 *P. viticola* samples, collected from wild and cultivated grapes in Northeast America, and from the main grape-growing regions in which grapevine downy mildew occurs. Using nuclear and mitochondrial gene sequences, we found that all invasive grapevine downy mildew populations worldwide belonged to a small clade of the species *P. viticola* f. sp. *aestivalis*, which parasitizes the *V. aestivalis* summer grape in Northeast America (Supplementary Photo S1). Using microsatellite markers on *P. viticola* f. sp. *aestivalis*, we found that there was little admixture between populations and that invasive populations had a low genetic diversity. The European populations had the highest level of diversity of all invasive populations and harbored all but one of the haplotypes present in other invaded areas. Using ABC scenario testing, we confirmed that *P. viticola* f. sp. *aestivalis* was first introduced into Western Europe, whence it spread to Central and Eastern Europe. The successful invasive populations in Europe then served as the source for secondary introductions into other grape-growing regions of the world, such as Northeast China, South Africa, and Australia. A third bridgehead effect occurred later, with the spread of the disease from Southeastern Australia to Argentina. All grapevine downy mildew invasions therefore stem from an initial single introduction event in Europe, followed by secondary and tertiary introductions via bridgehead effects. These introductions were probably mediated by the transfer of grapevine material by humans, as settlers imported European cultivars for the establishment of “New World” vineyards during the 19th century (*37*). Our findings of a strong bottleneck following the introduction into Europe and the common origin of all introduced populations worldwide provide essential knowledge for guiding breeding for resistance to grapevine downy mildew. The identification of these historical pathways also improves our understanding of biological pest and pathogen invasions.

## Results

### The cryptic species *Plasmopara viticola* f. sp. *aestivalis* is the origin of all invasive downy mildew populations worldwide

We reconstructed the genetic relationships between an extensive set of *P. viticola* strains collected from wild or cultivated grape species in native areas and areas of introduction, by sequencing DNA fragments from the mitochondrial cytochrome-*b* (*cytb, n* = 1,299), β-tubulin (*tub*, *n* = 424 samples) and ribosomal 28S (*r28S*, *n* = 536) genes (Fig. 1, Supplementary Fig. S1 and Table S1). The β-tubulin tree provided the highest resolution (Fig. 1A, with 56 distinct haplotypes versus 32 haplotypes for *cytb* and 9 haplotypes for *r28S*, Supplementary Fig. S2). Nevertheless, all the trees presented highly differentiated lineages with contrasting host ranges (Fig. 1A, Supplementary Fig. S2), consistent with the previously demonstrated existence of five cryptic species (*33*, *34*). Three *formae speciales* were found on cultivated grapes in North America, but only *P. viticola* f. sp. *aestivalis* was found in introduced populations worldwide (Fig. 1 and Supplementary Fig. S2). In North America, this *forma specialis* was found on *V. aestivalis* and on *V. labrusca sensu lato* (i.e. including *V. labrusca* and its main artificial hybrids), two *Vitis* species that have recently diverged (*38*).

**Figure 1.**
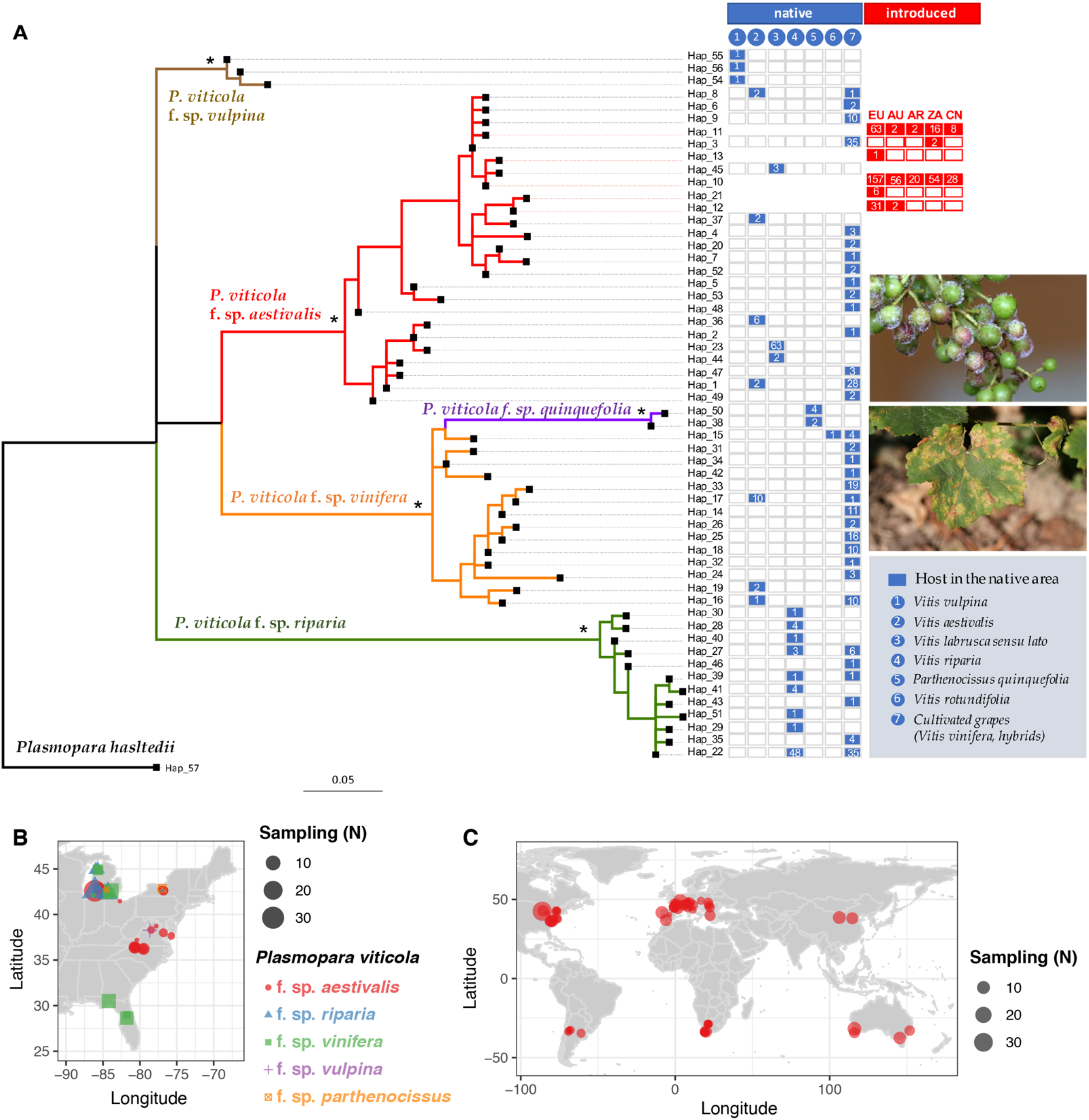
Phylogenetic relationships, sampling, and geographic distribution of *Plasmopara viticola* haplotypes of the β-tubulin (*tub*) gene. **A:** The maximum likelihood phylogenetic tree is rooted with the *Plasmopara hasltedii tub* sequence. Nodes with a star (*) are supported with a bootstrap value greater than 90%. The branches of the tree are color-coded according to the five *P. viticola formae speciales* (ff. spp.). Colored and empty boxes on the right side of the tree show the host plant on which the haplotype was found in the native area or its geographic location on the introduced areas. Geographic codes include Europe (EU), Australia (AU), Argentina (AR), South Africa (ZA) and China (CN). The number of isolates carrying a given haplotype is indicated by the number within the boxes. Photos on the right show the grapevine downy mildew pathogen, *P. viticola*, infecting young grape berries (top) and typical lesions on a leaf (bottom) of *Vitis vinifera* in Europe. **B:** Map showing the sample size (N) and geographic location of origin of the strains from the five *formae speciales* occurring in Northeast America. **C:** Distribution of the sampling sites of the *P. v*. f. sp. *aestivalis* strains across the main wine-producing regions worldwide.

### A host shift from *V. aestivalis* at the origin of the invasive downy mildew populations and population subdivision in invasive populations

We used highly polymorphic microsatellite markers to obtain a finer resolution of the population structure within *P. viticola* f. sp. *aestivalis*, to elucidate its worldwide population structure and invasion routes. We genotyped 1,974 diploid strains with eight microsatellite markers (Supplementary Table S2), which, together with the three sequenced fragments (Supplementary Table S1), revealed 1,383 distinct genotypes with this 11-marker dataset. We found that the native and introduced *P. viticola* f. sp. *aestivalis* populations were strongly differentiated, along the first axis of a discriminant analysis on principal components (DAPC) (*39*) (Supplementary Fig. S3). The second axis revealed a strong differentiation in North America between *P. viticola* f. sp. *aestivalis* populations collected on different *Vitis* species (Supplementary Fig. S3). Indeed, we identified specific genetic clusters on *V. aestivalis* (in yellow), *V. labrusca* (in black), and two differentiated clusters (brown and red) on the cultivated grape *V. vinifera* (Supplementary Fig. S3). The cluster on cultivated grapes (in brown) was genetically close to the cluster on the wild species *V. labrusca* (in black), probably indicating a host shift from *V. labrusca* in the native range. We inferred that the invasive populations originated from a host shift from *V. aestivalis*, as the population from this wild summer grape (yellow cluster) appeared genetically closest, and even overlapping in the DAPC, with the invasive populations (Supplementary Fig. S3A). However, the eight microsatellite markers had a low power to resolve the genetic structure between invasive populations (Supplementary Fig. S3C).

We therefore genotyped a subset of strains (*n* = 181) with 32 microsatellite markers, focusing on invasive populations and on the native *P. viticola* f. sp. *aestivalis* clusters that may have served as origin of the invasive populations (Supplementary Table S3, 174 unique genotypes). This 32-marker dataset confirmed the genetic patterns observed in the native range based on the 11-marker dataset, and, in particular, confirmed that the cluster on *V. aestivalis* (yellow) was the likely origin of all invasive populations worldwide (Fig. 2). This cluster was, again, the closest to all the invasive populations on the DAPC (Fig. 2D). Furthermore, using the 32-marker dataset, we showed that the cluster on *V. aestivalis* was the only cluster having common genetic ancestry with the invasive populations in the STRUCTURE Bayesian clustering analysis (*40*, *41*) (see the light blue, green or pink ancestry from *K* = 2 to *K* = 6 in genotypes collected from *V. aestivalis* on Fig. 2A). With the 32-marker dataset, we also showed that this was the closest cluster to all the invasive populations in the distance-based neighbor-joining tree (Fig. 2C). The 32-marker dataset further identified two genetic clusters on *V. aestivalis* in the native range (Fig. 2A): the yellow cluster identified above, genetically close to the invasive populations found on cultivated grapes worldwide (Fig. 2D), and the red genetic cluster, corresponding to strains collected on cultivated grapes in North America (Fig. 2A). This suggests the occurrence of two distinct host shifts from the wild summer grape *V. aestivalis* to cultivated grapes, one in the native range, from the red genetic cluster, and the other, from the yellow cluster, giving rise to all invasive populations worldwide.

**Figure 2.**
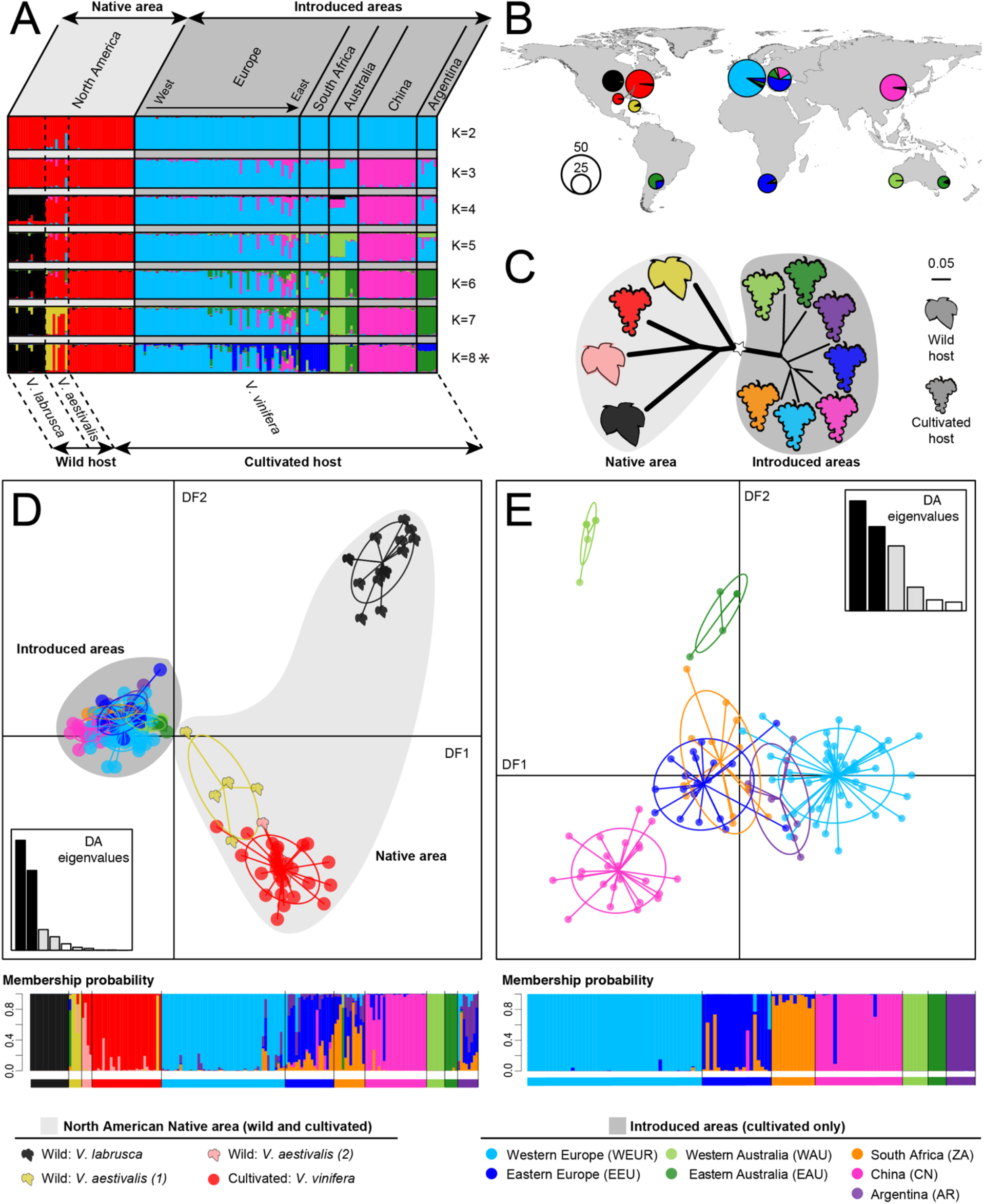
Worldwide population genetic structure of *Plasmopara viticola* inferred from the 32-microsatellite marker dataset. **A**: Estimated individual ancestry according to the Bayesian clustering approach of STRUCTURE, for two to eight clusters (*K*). Each individual is represented by a thin vertical line, partitioned into *K* colored segments representing the estimated genome ancestry fractions for each cluster. Individuals from different continents (labeled at the top of the figure) are separated by black continuous lines; strains from different host species (labeled at the bottom of the figure) are separated by black dotted lines; within these categories, individuals are sorted by increasing longitude. The figure shown for each *K* corresponds to the highest probability run from 10 replicates, with the best-fitting *K* value indicated by a star. **B**: STRUCTURE admixture proportions for samples averaged across populations for *K* = 8. Pie chart size is proportional to sample size. **C**: Neighbor-joining (NJ) tree based on the Cavalli-Sforza chord distance. Population branching with high bootstrap support (i.e., >75%), indicated by large widths. Wild and cultivated grapes are represented by different symbols. North America and other continents are separated into different gray circles. **D** and **E**: Scatterplot of the first two discriminant functions (DFs) from the discriminant analysis on principal components (DAPC), showing *P. viticola* individual genotypes for the full dataset (D) and for the subset excluding populations from the native North American range (E). The histogram insets of each DAPC scatter plot show the proportion of variance explained by each DF, according to their respective eigenvalues. The bar plots below each scatter plot represent the probability of the strain belonging to each group, based on all DFs of the DAPC. The populations studied are as follows: North American *P. viticola* strains collected on wild *V. labrusca*, *V. aestivalis* group 1 (yellow) and 2 (red); North American strains collected on cultivated *V. vinifera*. Strains from the rest of the world collected on cultivated *V. vinifera* including, strains from Western and Eastern Europe (WEUR and EEUR), Western and Eastern Australia (WAUS and EAUS), China (CN), South Africa (ZA), and Argentina (AR).

The 32-marker dataset also revealed genetic differentiation between invasive populations in the main wine-producing regions worldwide (Fig. 2). The two distinct genetic groups of *P. viticola* previously identified in Western and Eastern European vineyards (*22*), respectively, were also detected here in STRUCTURE clustering analyses (Fig. 2A) and with the DAPC (Fig. 2E). Furthermore, the populations in the different invaded areas also displayed significant differentiation, as shown by the significant *F_ST_* values (Supplementary Table S4 and S5), STRUCTURE analyses (Fig 2A) and the DAPC analyses focusing on invasive populations (Fig. 2E). We also found two differentiated populations in Australia (Fig. 2).

### Lower diversity in invasive *P. v*. f. sp. *aestivalis* populations

Genetic diversity in invasive *P. v*. f. sp. *aestivalis* populations worldwide was much lower than that in native populations. Only a few closely related haplotypes of the three sequenced DNA fragments were present in the invaded areas, whereas considerable diversity was detected in the native *P. v*. f. sp. *aestivalis* populations (Fig. 1, Supplementary Fig. S2, Table S1). The resulting low nucleotide (π) and haplotype (H) diversities in the invasive populations (Fig. 3A, Supplementary Table S1) indicated the occurrence of severe bottlenecks during the invasion process. European populations displayed a smaller decrease in genetic diversity than the populations of other invaded areas (Figs. 1 and 3A, Supplementary Fig. S2, Table S1), suggesting that the bottlenecks occurring in Europe were milder. Furthermore, all but one of the invasive haplotypes worldwide were present in Europe, for the three DNA fragments analyzed (Fig. 1 and Supplementary Fig. S2). Together, these findings suggest that Europe may have acted as a bridgehead for secondary invasions in the rest of the world.

**Figure 3.**
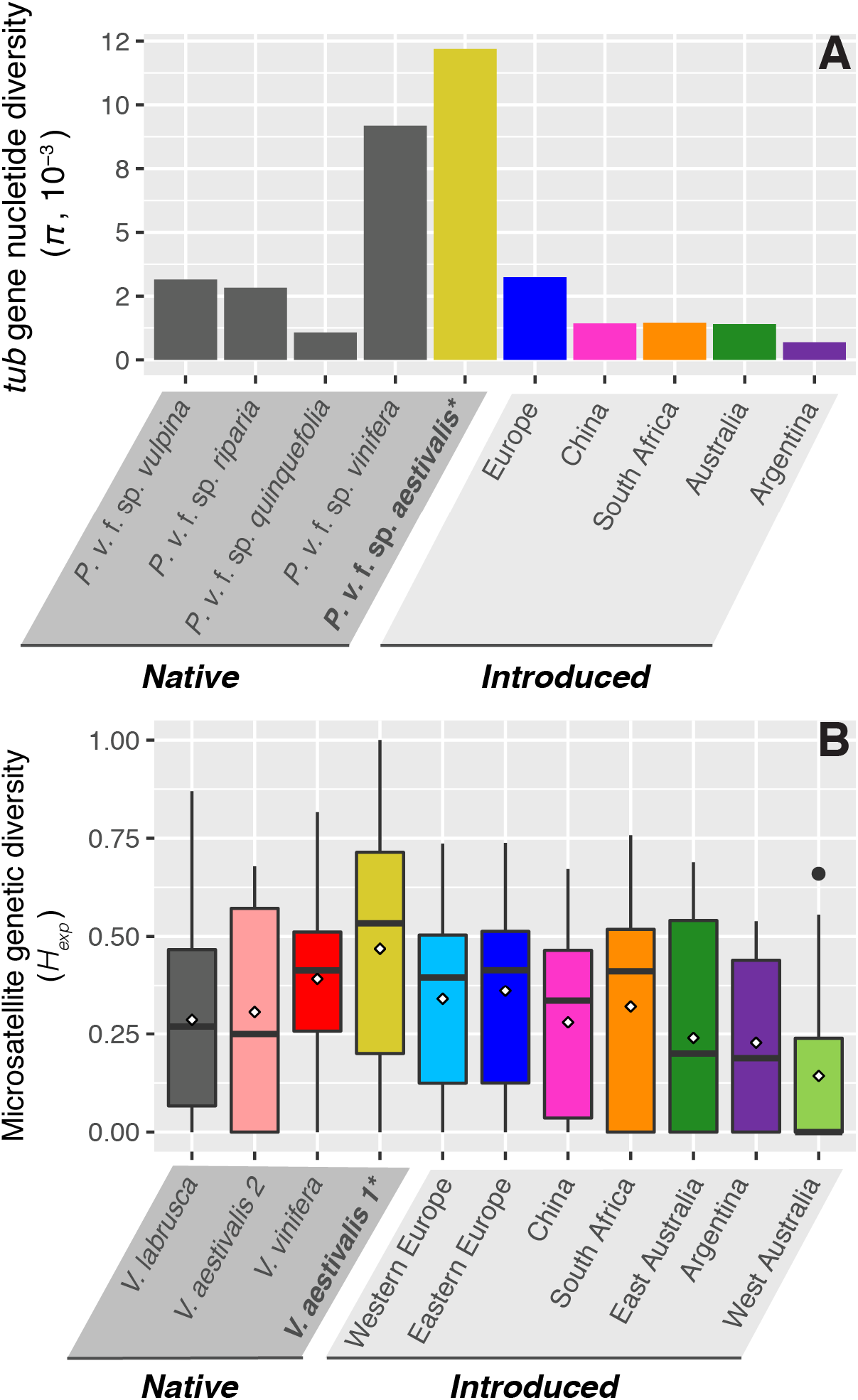
Lower diversity in invasive populations of *Plasmopara viticola forma speciale aestivalis*. **A**) Bar plot showing the nucleotide diversity (π, per site) of the β-tubulin (*tub*) gene sequence. **B**) Box plot showing the distribution of expected heterozygosity (*H_exp_*) for the 32 microsatellite markers.

Both microsatellite datasets confirmed the much lower levels of genetic diversity in invasive populations than in native populations, as shown by the compact clustering of invasive genotypes in the DAPC (Fig. 2D, Supplementary S3A), and by diversity indices (Fig. 3B, Supplementary Tables S2, S3, and S6 to S11). For example, private allelic richness was one order of magnitude lower in the invasive populations worldwide than in the native range. Microsatellite genetic diversity was also much lower in introduced than in native populations, with European populations having intermediate values (Fig. 3B, Supplementary Tables S2 and S3).

This pattern of diversity is, thus, consistent with the hypothesis that Europe acted as a bridgehead, with a first wave of invasion to Europe originating from the yellow *P. v*. f. sp. *aestivalis* cluster (Fig. 2 and Supplementary Fig. S3) occurring on the wild summer grape in the native Northern American range, and a second wave of invasions subsequently occurring from Europe to other vineyards throughout the world.

### Worldwide invasion history of *P. viticola* reconstructed by ABC-RF scenario testing

We formally compared the likelihoods of alternative invasion history scenarios involving the most likely population of origin in the native range (*i.e*., the yellow genetic cluster on *V. aestivalis*), as identified based on previous analyses, and all the invasive introduced populations, using the 32-microsatellite marker dataset and an approximate Bayesian computation random forest (ABC-RF) statistical framework (*42*). In ABC-RF, genetic data are simulated under different demographic scenarios, and summary statistics from the resulting simulated data are statistically compared with those obtained from the observed data (*43*–*45*). We used an iterative process to infer the various invasion events while keeping a tractable number of scenarios to be compared with the ABC-RF (*46*). We first identified the most likely demographic scenario(s) composed of bifurcation and admixture events considering the native populations and the invasive populations that corresponded to the most ancient introduction events (i.e., with dates of first disease records outside the native range between 1878 and 1889, Fig. 4B). We then considered the other invasive populations successively following increasing dates of first disease records and tested using ABC-RF what was their population of origin.

**Figure 4.**
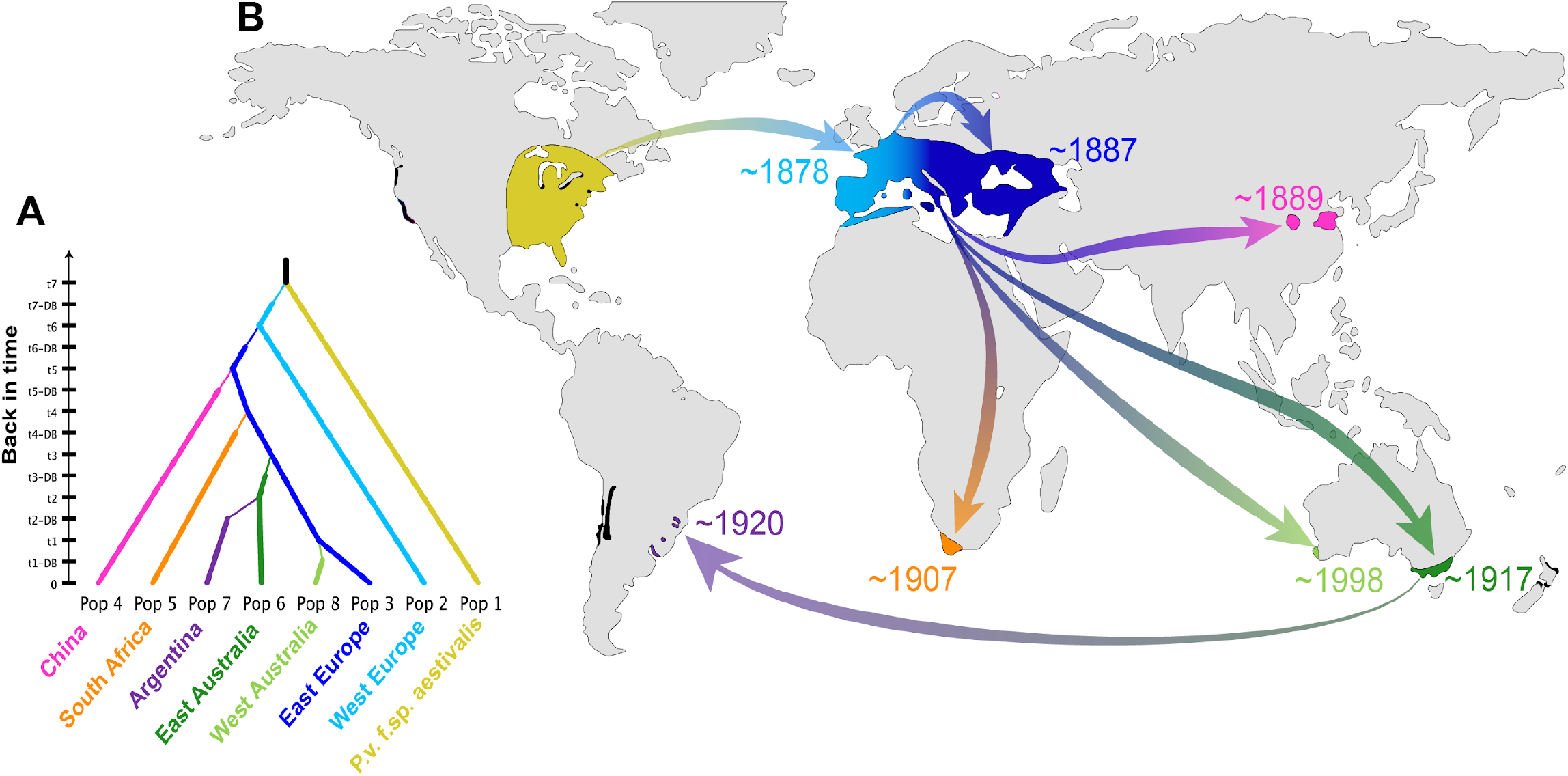
Worldwide invasion history of *Plasmopara viticola forma speciale aestivalis* inferred by the approximate Bayesian computation random forest (ABC-RF) approach. (**A**) The best population divergence scenario inferred by ABC-RF (Supplementary Table S12). (**B**) Geographic representation of the invasion scenario with the highest likelihood based on the ABC-RF analysis; areas are shown in color, based on their population assignment, as identified in Figure 2. Dates represent the first report of the disease in each area of introduction. Arrows indicate the most likely invasion pathways. Vineyards in other regions of the world not included in this study are indicated in black.

As a first step, we considered 18 scenarios of introduction from the most likely population-of-origin in the native range (the yellow cluster on *V. aestivalis*) and the first reported invasive populations from historical records (*i.e*., in Western and Eastern Europe, in ~1878 and ~1887, respectively, and in China, in ~1889; Supplementary Table S12 and Fig. S4). The most likely scenario identified (S11 in Supplementary Table S12 and Fig. S4) involved an initial introduction into Western European vineyards, followed by a spread to Eastern Europe (as previously shown (*22*)), and then an expansion to Eastern China from Eastern Europe. This scenario was the most strongly supported, with a posterior probability of 0.42 ± 0.01 (Supplementary Table S12).

The disease was later reported in South Africa in ~1907 and in eastern Australia in ~1917. For these two introduction events (steps 2 and 3 in Supplementary Table S12, Fig. S5 and S6), the ABC-RF again identified Eastern Europe as the most likely population of origin, with posterior probabilities of 0.78 ± 0.03 and 0.71 ± 0.02, respectively. The disease was then reported in Argentina in ~1920, and the ABC-RF analysis suggested that this introduction was probably from Eastern Australia, with a posterior probability of 0.70 ± 0.02 (step 4 in Table S1, Supplementary Fig. S7). The most recent first report of the disease was in Western Australia in ~1917, for which an Eastern Australian origin has been suggested (*28*). However, the ABC-RF statistical framework, based on a larger set of microsatellite markers and more extensive sampling of *P. viticola* populations worldwide, again identified Eastern Europe as the most likely population of origin for the pathogen in Western Australia, with a posterior probability of 0.73 (step 5 in Supplementary Table S12 and Fig. S8).

The most likely global invasion history is summarized in Figure 4. The classification error rate (i.e., prior error rate) for the first step was relatively high (38%), reflecting the difficulty distinguishing between a first introduction into Eastern or Western Europe, as these populations are genetically very close. By contrast, all subsequent steps in the ABC-RF analysis were well resolved, with a low prior error rates, ranging from 7% to 12% (Supplementary Table S12). The final global scenario (Fig. 4) was strongly supported by the genetic data, as shown by the low prior error rate of 7% (Supplementary Table S12). Posterior model checking and goodness-of-fit assessment of the model-posterior distributions showed that the inferred global invasion scenario generated genetic summary statistics consistent with the observed data, providing high confidence in the inferred scenario. The 10,000 simulations from posterior parameter distributions under the best model produced summary statistics very close to those obtained from the observed dataset (Supplementary Fig. S9), with only 24 of 256 statistics falling in the tail of the predictive probability distribution of statistics calculated from the posterior simulations (i.e., *P* < 0.05 or *P* < 0.95). Moreover, none of the *P-values* remained significant after correction for multiple comparisons (*47*). The inferred invasion scenario (Fig. 4) thus fitted the observed genetic data well.

## Discussion

Using extensive sampling and a powerful statistical framework, we have elucidated the worldwide invasion history of grapevine downy mildew. We show here that a specific genetic cluster of *P. v*. f. sp. *aestivalis*, which parasitizes the wild summer grape *V. aestivalis* in the native range, was the source of the invasion responsible for a devastating pandemic on cultivated grapevines in Europe at the end of the 19th century (*9*, *12*). Severe bottlenecks occurred, with very few haplotypes from the native lineage introduced into Europe. In the inferred invasion scenario (Fig. 4), Europe then served as a bridgehead for a second wave of invasions worldwide, spreading the disease further afield, to Eastern China, South Africa and Eastern Australia. A third wave of expansion probably occurred from Eastern Australia to Argentina. The much more recent introduction in Western Australia (1998) also seems to have originated from Europe, raising questions about the efficacy of quarantine regulations. The contribution of European populations to grapevine downy mildew invasions worldwide reflects the key role of Europe in the trading of plant material during the development of “New World” viticulture. The phylloxera crisis intensified the importation of plant material from North America to Europe and from Europe to “New World” vineyards (*48*). A role for Europe as a hub for the invasion of “New World” vineyards by grapevine pathogens has also been suggested for phylloxera and for the soil-born nematode vector of the fanleaf degeneration virus (*48*, *49*).

Controlling the introduction of pests and diseases is one of the greatest challenges in viticulture, with important economic and environmental consequences. The genetic relationships and diversity inferred here could be used to guide quarantine regulations for grape-growing countries, to prevent further invasions. Introducing new strains or lineages of *P. viticola* would significantly increase the diversity of invasive populations and create opportunities for genetic admixture. Indeed, we found that, in North America, the cultivated grape (*V. vinifera*) was parasitized by a *P. viticola* lineage (f. sp. *vinifera*) other than the one identified here as invasive (f. sp. *aestivalis*). The *P. viticola* f. sp. *vinifera* lineage has not yet been found on other continents, but could also become invasive should it be introduced. An increase in genetic variation through migration and/or admixture would probably facilitate the adaptation of *P. viticola* to pesticides and resistant cultivars (*50*). New invasions by additional *P. viticola* strains or lineages would have a major effect on the wine industry, by destabilizing grapevine protection and canceling out the sustained efforts of breeders to obtain good-quality grapevine varieties resistant to downy mildew. The rich historical records available for grape diseases make these diseases excellent case studies for obtaining fundamental insight into the processes underlying biological invasions, showing that worldwide biological invasions can result from the secondary dispersal of a particularly successful invasive population, as for grape phylloxera (*48*, *51*). Our findings show that even pathogens subject to very strong bottlenecks and without admixture as a means of generating diversity can achieve successful invasions worldwide, by retaining an ability to evolve rapidly, and thus to develop new virulence against plant resistance (*52*–*54*) and resistance to fungicides (*21*, *55*, *56*).

## Materials and methods

### *Plasmopara viticola* sample collection and DNA extraction

*Plasmopara viticola* isolates were collected as sporulating lesions from 163 sites in the native (North America) or invasive (Europe, China, South Africa, Australia, and Argentina; Fig. 1B and 1C, Supplementary Fig. S1; Table S1, S2 and S3) range. This sampling covers the main grape-growing regions in which the climate is favorable for the disease. In Northeast America, samples were collected from five wild *Vitis* species (*V. riparia, V. labrusca*, *V. aestivalis, V. vulpina* and *P. quinquefolia*), and cultivated grapevines (*V. vinifera* and interspecific hybrids). In invaded areas, all samples were collected from diseased *V. vinifera* cultivars. Sampling was performed as previously described (*22*). Oil spots were freeze-dried overnight, and DNA was extracted with standard CTAB and phenolchloroform methods (*55*, *57*).

### DNA sequencing and analysis

For the identification of cryptic *P. viticola* species, three DNA fragments, from the 28S ribosomal RNA (*r28S*), *β*-tubulin (*tub*) and mitochondrial cytochrome-*b* (*cytb*) genes, were amplified by polymerase chain reaction (PCR) and sequenced (Supplementary Table S1). DNA amplification, sequencing and assembly were performed as described by Rouxel *et al*. (*34*) for *r28S* and *tub*, and as described by Chen *et al*. (*55*) and Giresse *et al*. (*58*) for *cytb*. Sanger sequencing was performed at the Genoscope (French national sequencing center, Evry, France). For each DNA fragment, sequences were aligned with *Muscle* (*59*) implemented in Seaview v.4.6.2 (*60*). Final alignment lengths after cleaning were 499 base pairs (bp) for *tub*, 685 bp for *cytb* and 703 bp for *r28S*. Distinct haplotypes were identified with DnaSP v5.10.1 (*61*). For each sequence alignment, genetic relationships between haplotypes were inferred with a maximum likelihood (ML) method and a GTR substitution model implemented in PhyML v.3.0 (*62*). Node support was estimated by calculating 1,000 bootstraps. Each phylogeny was rooted with *Plasmopara hasltedii* sequences obtained from a recent genome assembly (*63*). Nucleotide (π, per site) and haplotype (H) genetic diversities were estimated with the DnaSP program.

### Microsatellite genotyping

We constructed two different microsatellite datasets for analysis of the genetic structure and diversity of *P. viticola* f. sp. *aestivalis* populations across the main vineyards worldwide, identification of the most likely host-of-origin, and elucidation of the invasion routes followed. The first dataset included an extensive sampling of 1,974 strains genotyped for eight microsatellite loci (ISA, Pv7, Pv13, Pv14, Pv16, Pv17, Pv31 and Pv39) (*23*, *57*) (Supplementary Table S2). Microsatellite PCR amplification and genotyping were conducted as previously described (*57*). The repeatability of genotype scoring was checked by genotyping 5% of the samples twice. For the analyses, genotypes were built by combining the eight microsatellite markers and the three DNA sequences. Genotypes with more than 60% missing data were discarded. A single genotype was retained when putative clone-mates (i.e., repeated multilocus genotypes or MLGs within a given vineyard) were identified. The final first genotype dataset encompassed 1,314 MLGs for the eight microsatellite markers and 1,383 MLGs for the 11 markers (combining the eight microsatellite markers with haplotypes for the three DNA fragments) from 105 localities (Supplementary Table S1).

The second dataset consisted of 181 samples genotyped for 34 microsatellite markers: ISA, Pv7, Pv14, Pv16, Pv17, Pv39, Pv65, Pv67, Pv74, Pv76, Pv83, Pv87, Pv88, Pv91, Pv93, Pv101, Pv103, Pv104, Pv124, Pv126, Pv127, Pv133, Pv134, Pv135, Pv137, Pv138, Pv139, Pv140, Pv141, Pv142, Pv143, PvEST2, PvEST9 and PvEST10 (*35*, *57*). The repeatability of genotype scoring was checked by genotyping 5% of the samples twice. The mean error rate was ≤ 0.020 ± 0.005. Genotypes with missing data for more than 16 markers (≥ 50%) were discarded. Significant linkage disequilibrium (LD) was detected between two pairs of microsatellite markers (Pv87 and Pv104; Pv124 and Pv133) with a permutation test (1,000 permutations) implemented in GENEPOP v.4.2 (*64*). A single marker for each pair was therefore retained for data analyses (Pv104 and Pv124). After filtering for missing data and LD, the final dataset included 174 MLGs for 32 markers from 35 sites (Supplementary Table S3).

### Inference of population genetic structure and diversity

We used the two datasets to investigate genetic structure: one with 11 markers (combining the eight microsatellite markers with haplotypes for the three DNA fragments), and the other with 32 microsatellite markers. We ran similar analyses on the two datasets. We used two complementary individual-based methods to estimate the population structure: the Bayesian model-based clustering method implemented in STRUCTURE v.2.3.4 (*40*, *41*) and a discriminant analysis on PCA (DAPC) (*39*). We ran STRUCTURE with an admixture model with correlated allele frequencies, and uniform priors for the individual cluster of origin (*40*, *41*). We performed simulations with a number of putative clusters (*K*) ranging from 1 to 10. For each *K* value, we conducted 10 independent replicates and checked for convergence. Each analysis included a burn-in period of 50,000 Markov chain Monte Carlo (MCMC) iterations followed by 500,000 MCMC iterations for the estimation of model parameters. We determined the most relevant number of clusters (*K*) using (i) the log likelihood of the data for each *K* value (*40*), (ii) the rate of change of the log likelihood of the data with increasing *K* (*65*), and (iii) the visual inspection of clusters newly generated with increases in *K* (*66*). We used STRUCTURE HARVESTER v.0.6.94 (*67*) to visualize the likelihood and its rate of change across *K* values and replicated runs. For the identification of potentially different clustering solutions, results were summarized and displayed with CLUMPAK v.1.1 (*68*). This analysis was performed on the two datasets combining native and introduced populations, and also on each area separately, to explore finer genetic structure.

DAPC(*39*) does not make any specific assumptions about mating system or mode of reproduction. It provides a visualization of the genetic structure complementary to that provided by STRUCTURE, by summarizing the variance in allele frequencies summarized using principal components (PCs) and partitioning it between populations relative to the within-group variance. This analysis was performed on the two datasets, and on the native and introduced populations separately, to explore the finer genetic structure in the two areas. We used the ADEGENET v.2.0.1 R package (*69*) for conducting the DAPC. In accordance with the recommendations of the user guide, missing data were replaced by the mean value, the number of PCs used in the analysis was identified by the alpha-score optimization procedure and was set to 20 and 10 for the 11- and 32-marker datasets, respectively. *A priori* groups were defined according to host plant or continent, and further partitioning was implemented in accordance with the findings of STRUCTURE or previous studies (e.g. split between Eastern and Western Europe (*22*)). We assessed how distinct or admixed the clusters of the DAPC were, using scatter plots along the top discriminant axes (DA) and bar plots of individual membership probabilities for each group.

Genetic relationships between clusters were estimated with NJ population trees based on pairwise Cavalli-Sforza chord genetic distances (*70*) between populations computed with the program POPULATIONS v. 1.2.32 (*71*). The node supports were estimated with 1,000 bootstraps over loci. The trees were drawn with FigTree v.1.4.4 (*72*).

Allelic frequency differentiation between clusters was estimated with the Weir and Cockerham *F_ST_* estimator (*73*) implemented in FSTAT v.2.9.4 (*74*). The significance of pairwise *F_ST_* values was assessed with 100 random data permutations in FSTAT. We assessed the levels of microsatellite genetic diversity within clusters, using allelic richness (*A_r_*) and private allelic richness (*P_Ar_*), both standardized by a rarefaction method (*75*), to account for differences in sample size. We also calculated observed and expected heterozygosities (*H_o_* and *H_e_*, respectively), and the *F_IS_* fixation index (*73*). These statistics were calculated with ADZE v.1.0 (*76*), GENETIX v.4.05.2 (*77*), and FSTAT. Wilcoxon signed-rank tests were performed in the R statistical environment (*78*), to assess the significance of differences in genetic diversity between clusters.

### ABC-RF-based inferences of global invasion history

We reconstructed the worldwide invasion history of *P. viticola* f. sp. *aestivalis* using an approximate Bayesian computation (ABC) (*43*–*45*) random forest (RF) analysis (*42*, *79*, *80*). This ABC-RF analysis used the 32-microsatellite dataset and included eight populations: the seven genetic clusters identified outside the native range on the basis of the 32-microsatellite dataset and the wild population sampled in the native range on the summer grape *V. aestivalis*, which was the population genetically closest to the invasive clusters (Fig. 2). ABC-RF can estimate posterior probabilities of historical scenarios, based on historical data and massive coalescent simulations of genetic data. We used historical information (i.e., dates of first observation of invasive populations, Fig. 4) to define five sets of competing introduction scenarios, which were analyzed sequentially (Supplementary Table S12, Fig. S4 to S8). Step-by-step, each analysis made use of the results of the previous analyses, until the most recent invasive populations were considered. The scenarios for each analysis are detailed in Supplementary Table S12 and Figs. S4 to S8.

The scenario parameters (i.e., effective population size *N*, effective number of founders *NF*, admixture rate *R_a_*, duration of the bottleneck *DB*, and time of population split and admixture *T*) were considered as random variables drawn from prior distributions (Supplementary Table S13). We assumed a generalized stepwise mutation process with possible single-nucleotide insertion, to model a realistic mutation process of microsatellite loci in the coalescent simulations (*81*). We used DIYABC v.2.1.0 (*82*) to simulate genetic data for ABC-RF analyses. Simulated and observed datasets were summarized using the whole set of summary statistics proposed by DIYABC for microsatellite markers, describing the genetic variation for each population (e.g., mean number of alleles per locus, and mean genetic diversity), pair of populations (e.g., pairwise genetic diversity, mean *F_ST_* across loci between two populations, and shared allele distance), or trio of populations (e.g., maximum likelihood admixture estimates) (see the full list and details of the summary statistics in Supplementary Table S14). Linear discriminant analysis (LDA) components were also used as additional summary statistics (*83*). The total number of summary statistics ranged from 70 to 256, depending on the analysis (Supplementary Table S12).

We used the random forest (RF) classification procedure to compare the likelihood of the competing scenarios at each step with the R package *abcrf* v1.8.1 (*42*). RF is a machine-learning algorithm that uses hundreds of bootstrapped decision trees to perform classification, using the summary statistics as a set of predictor variables. Some simulations are not used in tree building at each bootstrap (i.e., the out-of-bag simulations), and are used to compute the “prior error rate,” which provides a direct method for estimating the cross-validation error rate. We built a training set ranging from 10,000 to 50,000 simulated microsatellite datasets for each scenario, with the same number of loci and individuals as the observed dataset (Supplementary Table S12). We then grew a classification forest of 500 or 1,000 trees based on the simulated training datasets. The size of the training set and number of decision trees was increased until the results converged over ten independent replicated RF analyses. The RF computation applied to the observed dataset provides a classification vote (i.e., the number of times a model is selected from the decision trees). We selected the scenario with the highest classification vote as the most likely scenario, and we estimated its posterior probability (*42*). We assessed the global performance of our chosen ABC-RF scenario, by calculating the prior error rate based on the available out-of-bag simulations and we repeated the RF analysis 10 times to ensure that the results converged.

We then performed a posterior model checking analysis on the final scenario, including all eight populations, to determine whether this scenario matched the observed genetic data well. Briefly, if a model fits the observed data correctly, then data simulated under this model with parameters drawn from their posterior distribution should be close to the observed data (*84*). The lack of fit of the model to the data with respect to the posterior predictive distribution can be measured by determining the frequency at which the observed summary statistics are extreme with respect to the simulated summary statistics distribution (hence, defining a tail-area probability, or *P*-value, for each summary statistic). We simulated 100,000 datasets under the full final scenario (256 summary statistics), and obtained a “posterior sample” of 10,000 values of the posterior distributions of parameters through a rejection step based on Euclidean distances and a local regression post-treatment (*43*). We simulated 10,000 new datasets with parameter values drawn from this “posterior sample,” and each observed summary statistic was compared with the distribution of the 10,000 simulated test statistics; its *P*-value was computed, and corrected for multiple comparisons (*47*). The simulation steps, the computation of summary statistics, and the model checking analysis were performed in DIYABC v2.1.0. All scenario comparisons were carried out in R, with the *abcrf* v1.8.1 package (*42*)

## Data availability

Unique haplotypes from the β-tubulin, cytochrome *b*, and ribosomal 28S sequence data were deposited in NCBI-Genbank under the accession codes: [to be announced, TBA]. The two microsatellite datasets and the sequence alignments are available via the INRAE Grapevine Downy Mildew Genomics Dataverse (https://data.inra.fr/dataverse/gdmg; doi: [TBA]).

## ACKNOWLEDGMENTS

We would like to thank all the people involved in *P. viticola* sampling: Anton Baudoin (Virginia Tech, USA), Odile Carisse (Agriculture and Agri-Food, Canada), David Gadoury (Cornell University, USA), Mizuho Nita (Virginia Tech, USA), Annemiek Schilder (University of California, Davis, USA), Andrew S. Taylor (DPI, Australia), Davide Gobbin (ETH, Switzerland), Marie-Laure Panon (Comité Champagne, France), Patrice Retaud (SRPV, France), Marie-Pascale Latorse (Bayer Crop-Science, France), Jorge Silva (Bayer CropScience, Portugal), Hanns Kassemeyer (Staatliches Weinbauinstitut Freiburg, Germany), Pal Kozma (University of Pécs, Hungary), Mauro Jermini (Agroscope, Switzerland), Sara Legler (UCSC, Italy), Atanas Atanassov (JGC, Bulgaria), Pere Mestre (INRAE, France), Jiang Lu (Shanghai JiaoTong University). This work was funded by ANR grant 07-BDIV-003 (*Emerfundis* project). YD received a grant from the French National Research Agency [GANDALF project, ANR-12-ADAP-0009] and MCF received a postdoctoral grant from the *Région Ile-de-France*. We thank the Center for Information Technology of the University of Groningen, and, in particular, Bob Dröge and Fokke Dijkstra, for their continuous support and for providing access to the *Peregrine* high-performance computing cluster.

