## Supplementary for "Europe as a bridgehead in the worldwide invasion history of grapevine downy mildew, *Plasmopara viticola*"

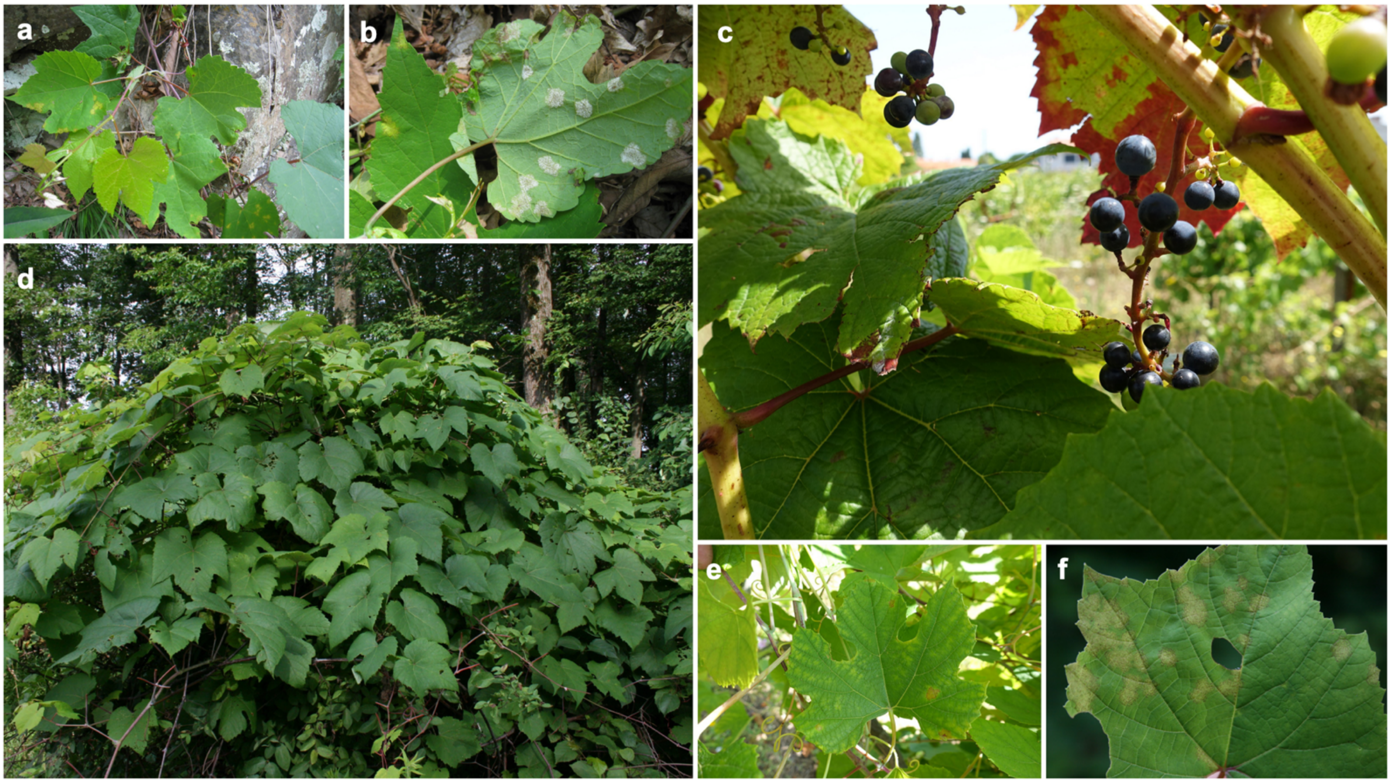


**Photo S1.** Grapevine downy mildew pathogen, *Plasmopara viticola*, on wild *Vitis aestivalis.*

a, b, e, f : typical pale yellow lesions (oil spots) on a diseased *V. aestivalis* leaf at spring.

c : cluster of the wild plant

d : infected plant on the edge of the forest

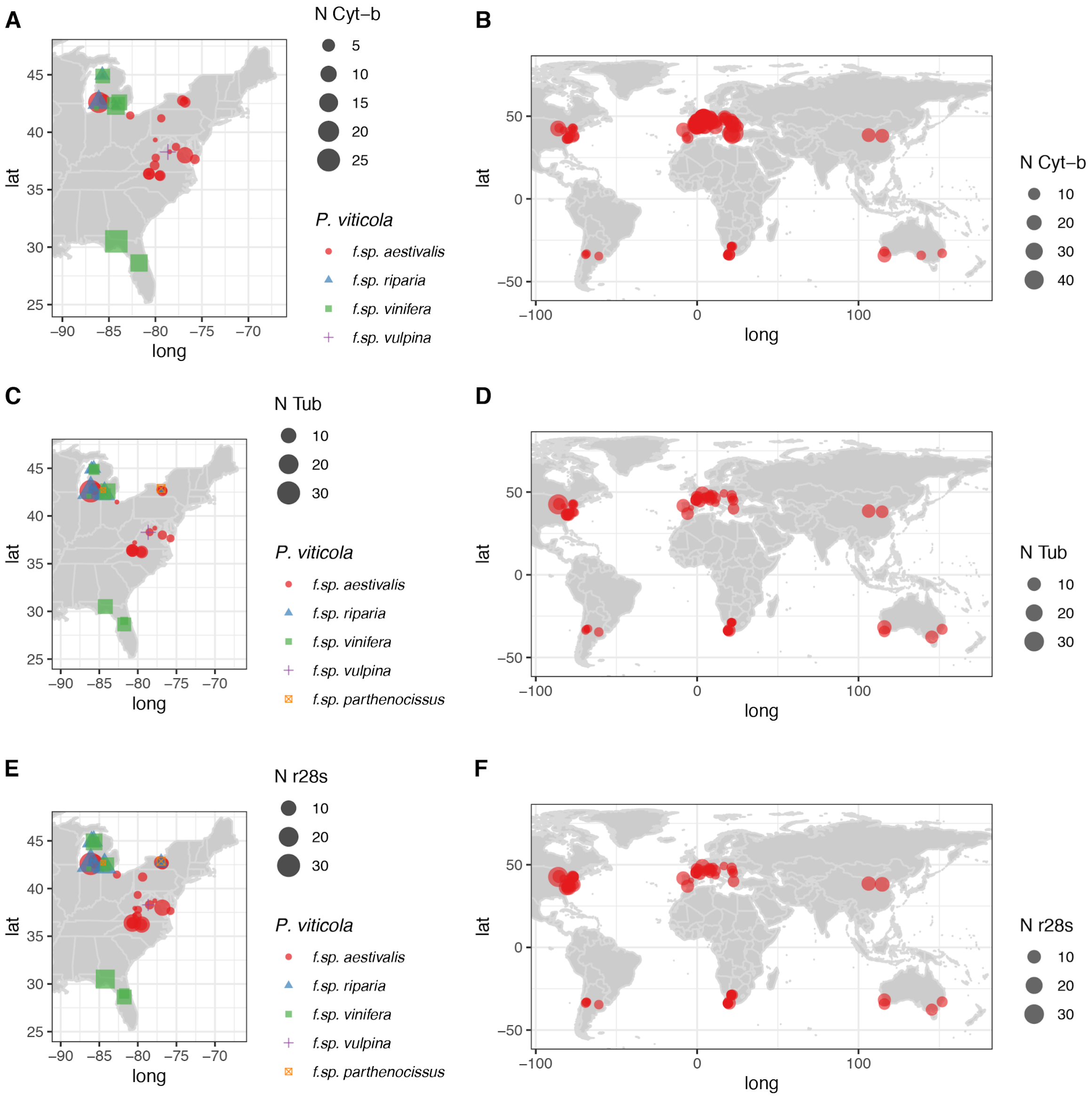


**Fig. S1.** Location, *formae speciales* and number of samples collected for the three genes sequenced in this study: (**A**, **B**) cytochrome-*b* (*cytb*, n = 1,294), (**C**, **D**) $\beta$-tubulin (*tub*, n = 420), and (**E**, **F**) ribosomal 28S (*r28S*, n = 534). The size of the point is proportional to the sample size. The shape and color of the points refer to the *P. viticola* formae speciales (f. sp.).

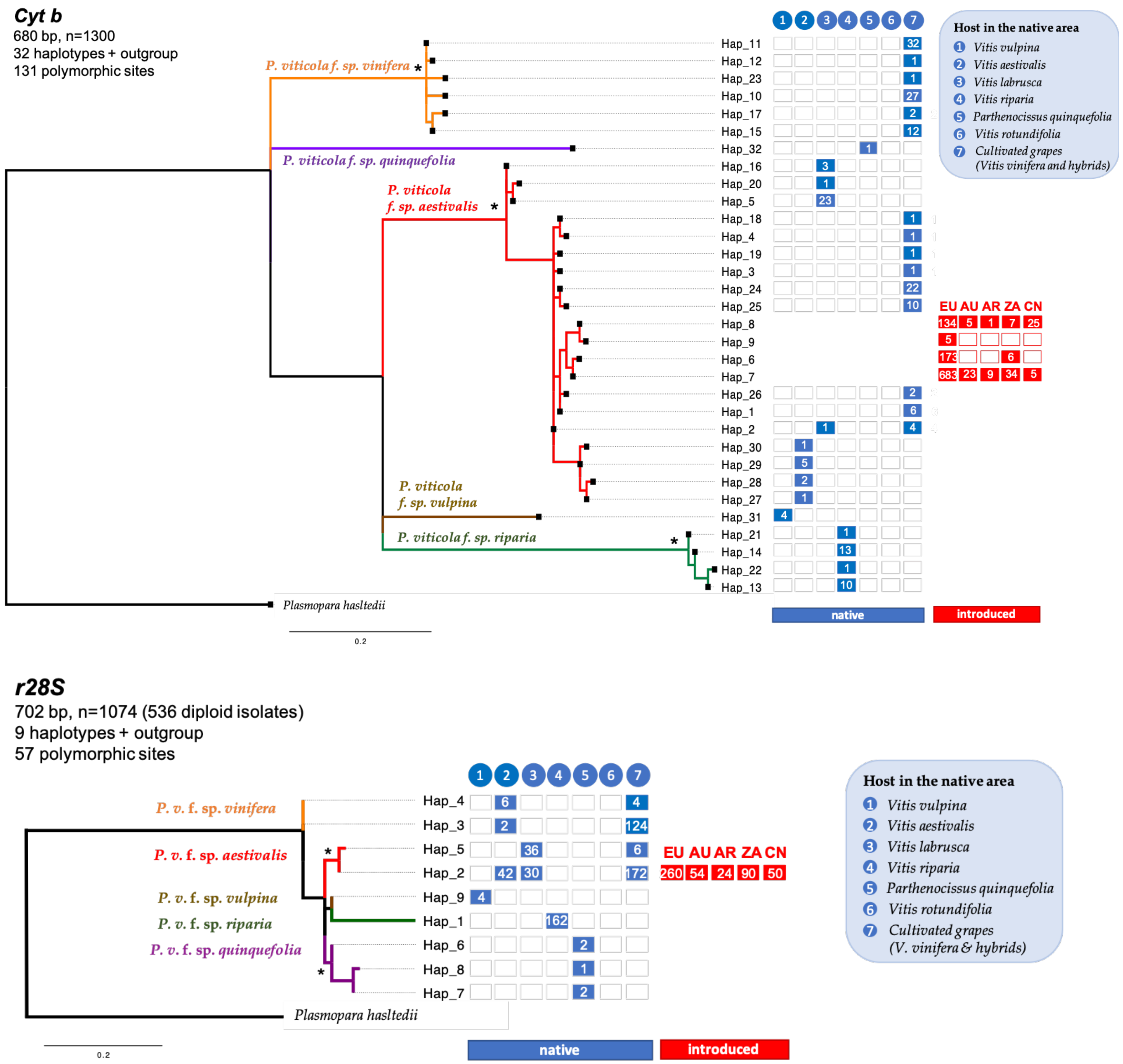


**Fig. S2**. Maximum likelihood phylogenetic relationships of *Plasmopara viticola* haplotypes for the mitochondrial *cytb* gene (top) and nuclear ribosomal *28S* (*r28S*, bottom). The tree is rooted with *Plasmopara hasltedii* sequences. Nodes with a star (*) are supported with a bootstrap value greater than 90%. The branches of the tree are color-coded according to the five *P. viticola formae speciales* (ff. spp.). Colored and empty boxes on the right side of the tree show the host plant on which the haplotype was found in the native area or its geographic location on the introduced areas. Geographic codes include Europe (EU), Australia (AU), Argentina (AR), South Africa (ZA) and China (CN). The number of isolates carrying a given haplotype is indicated by the number within the boxes.

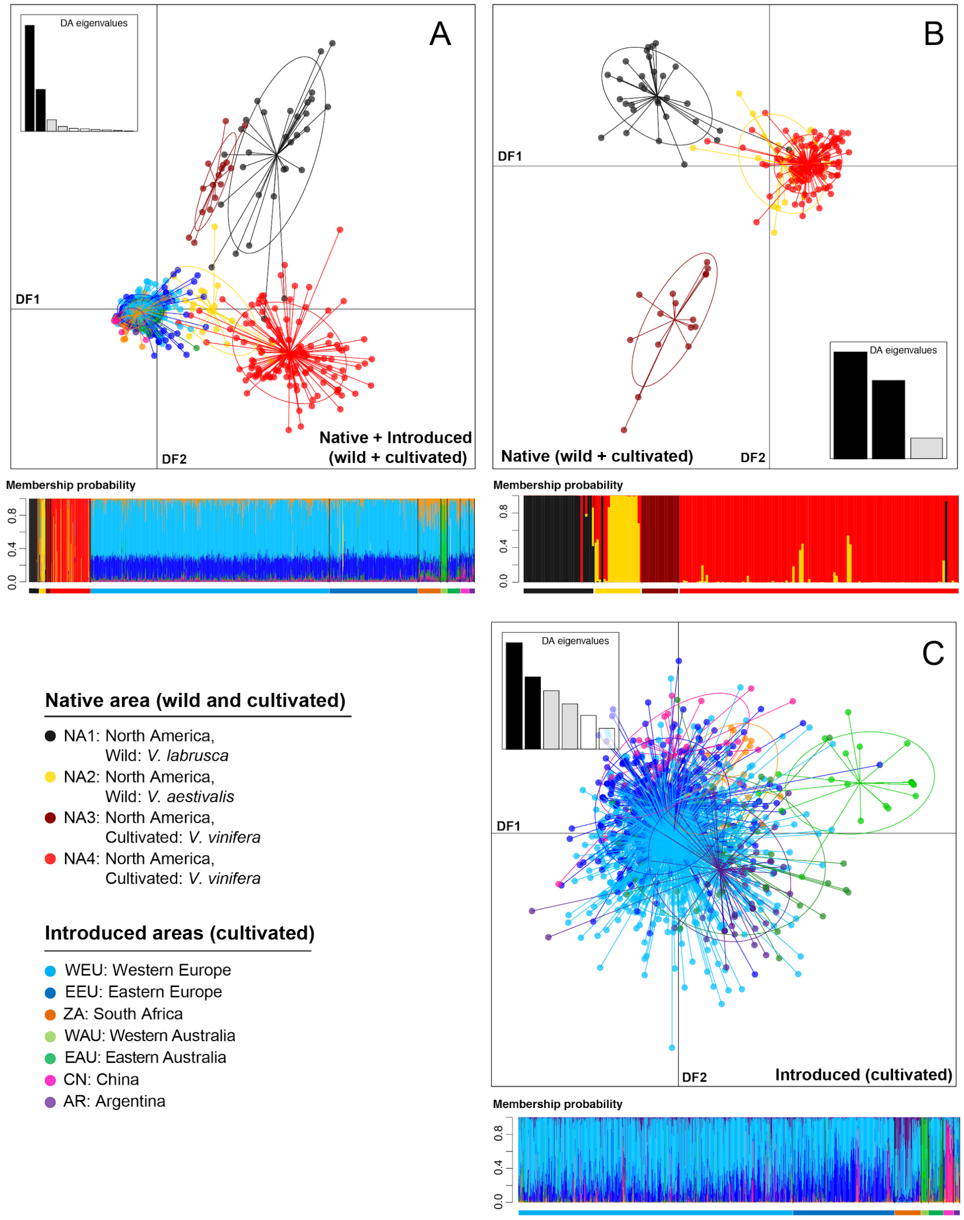


**Figure S3.** Worldwide population genetic structure of the downy mildew *Plasmopara viticola forma specialis aestivalis*, inferred from a discriminant analysis of principal components (DAPC) using genotypes of 1,383 strains based on eight microsatellite markers and sequences from three genes (cytochrome-*b*, β-tubulin, and ribosomal 28S). DAPC scatterplots for the first two discriminant functions (DF1 and DF2) show the clustering of the genotypes for the whole dataset (**A**), a subset focusing on the strains collected in the native North American range on wild and cultivated plants (**B**), and another subset in the introduced range on cultivated grapes (**C**). The inset in each panel shows the eigenvalues of the DFs. The barplot below each scatterplot displays the membership probabilities estimated from the DAPC for each genotype (vertical bars) to *a priori* groups defined by the host plant and locality, as indicated by the bars below the barplot, with the color indicating the country of collection, as indicated in the legend.

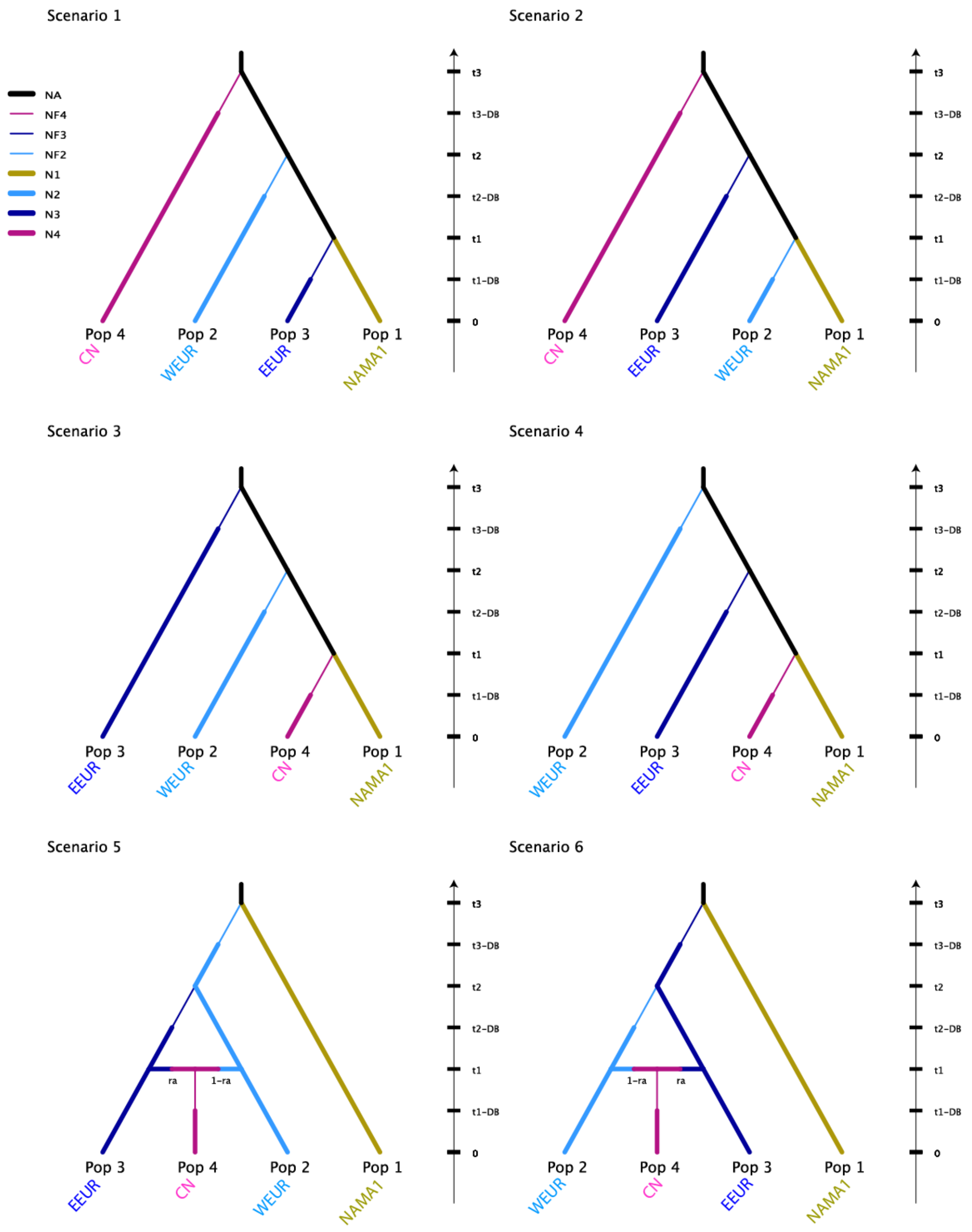


**Fig. S4.** Schematic representation of the scenarios tested in the step 1 of the ABC analyses to decipher the worldwide invasion routes of *Plasmopara viticola forma specialis aestivalis* populations. It includes 18 possible scenarios describing the evolution of four populations: *P. v.* f. sp. *aestivalis* source population sampled on *V. aestivalis* in Northeast America (NAMA1, yellow cluster in Fig. 2), the Western (WEUR) and Eastern European (EEUR) and Chinese (CN) clusters. For each analysis, the name of the most likely scenario is underlined and marked with an asterisk. Thin lines indicate bottlenecks. For parameter descriptions and priors see Table S12 and S13. Time is not to scale. (1/3)

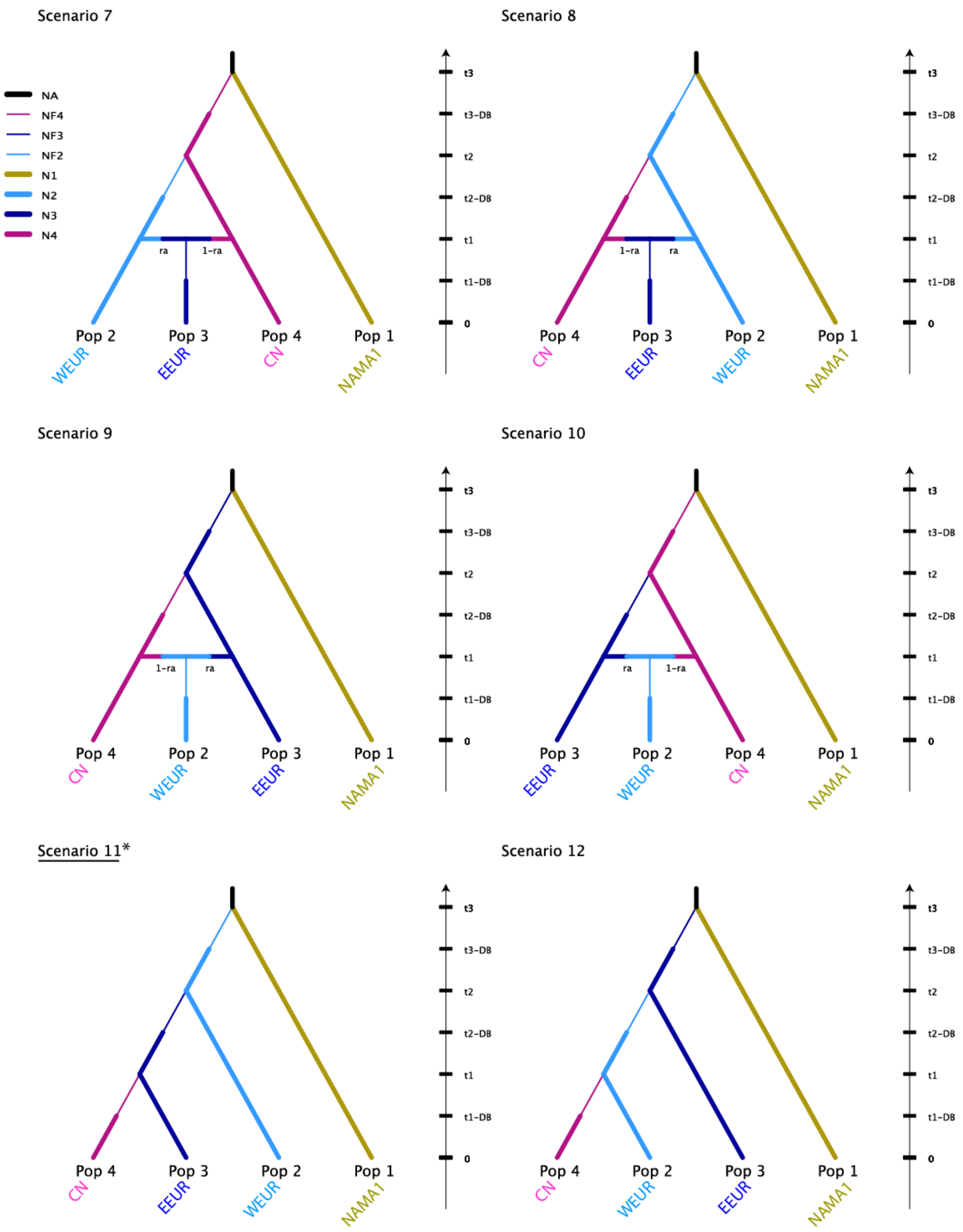


**Fig. S4.** Continued. (2/3)

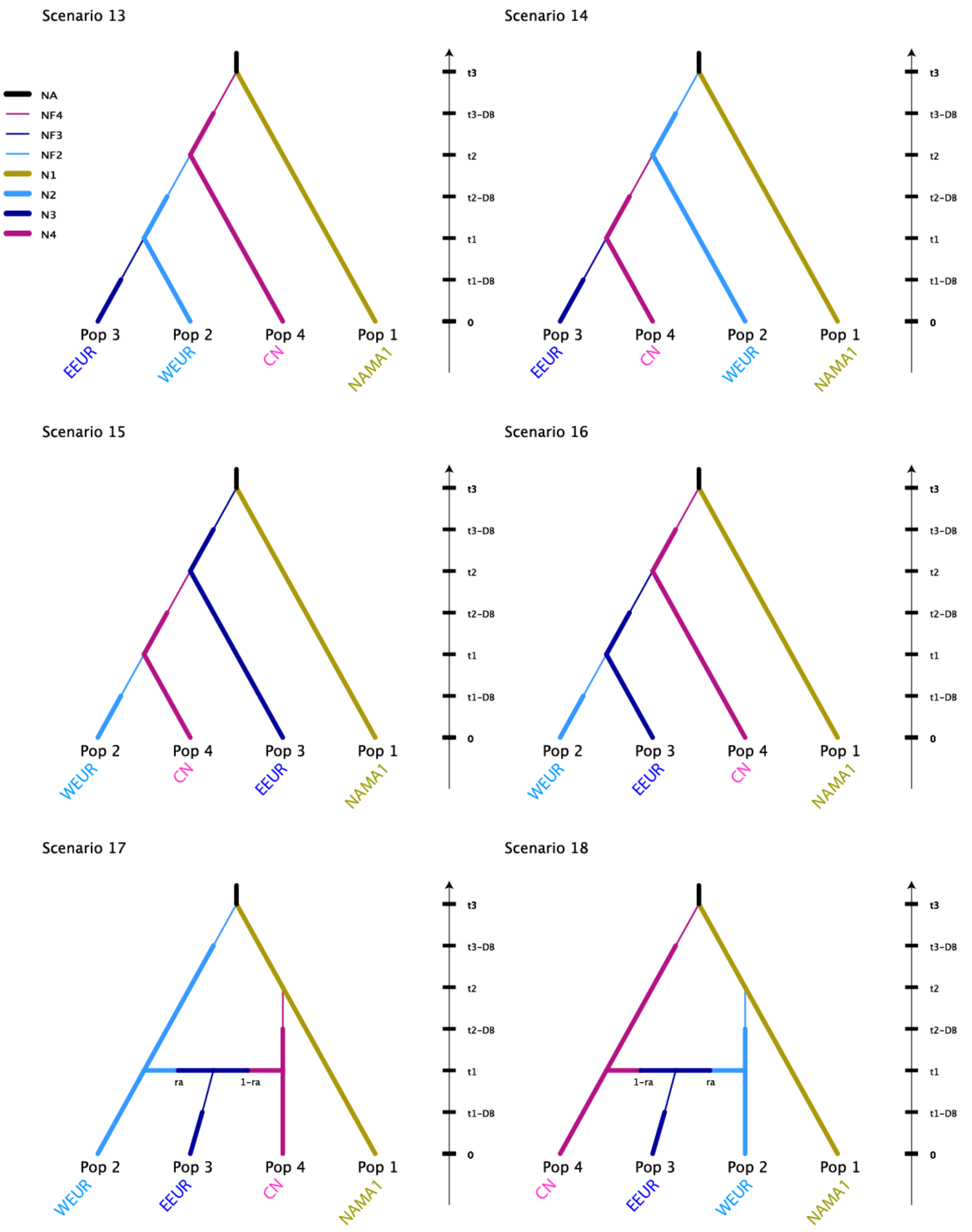


**Fig. S4.** Continued. (3/3)

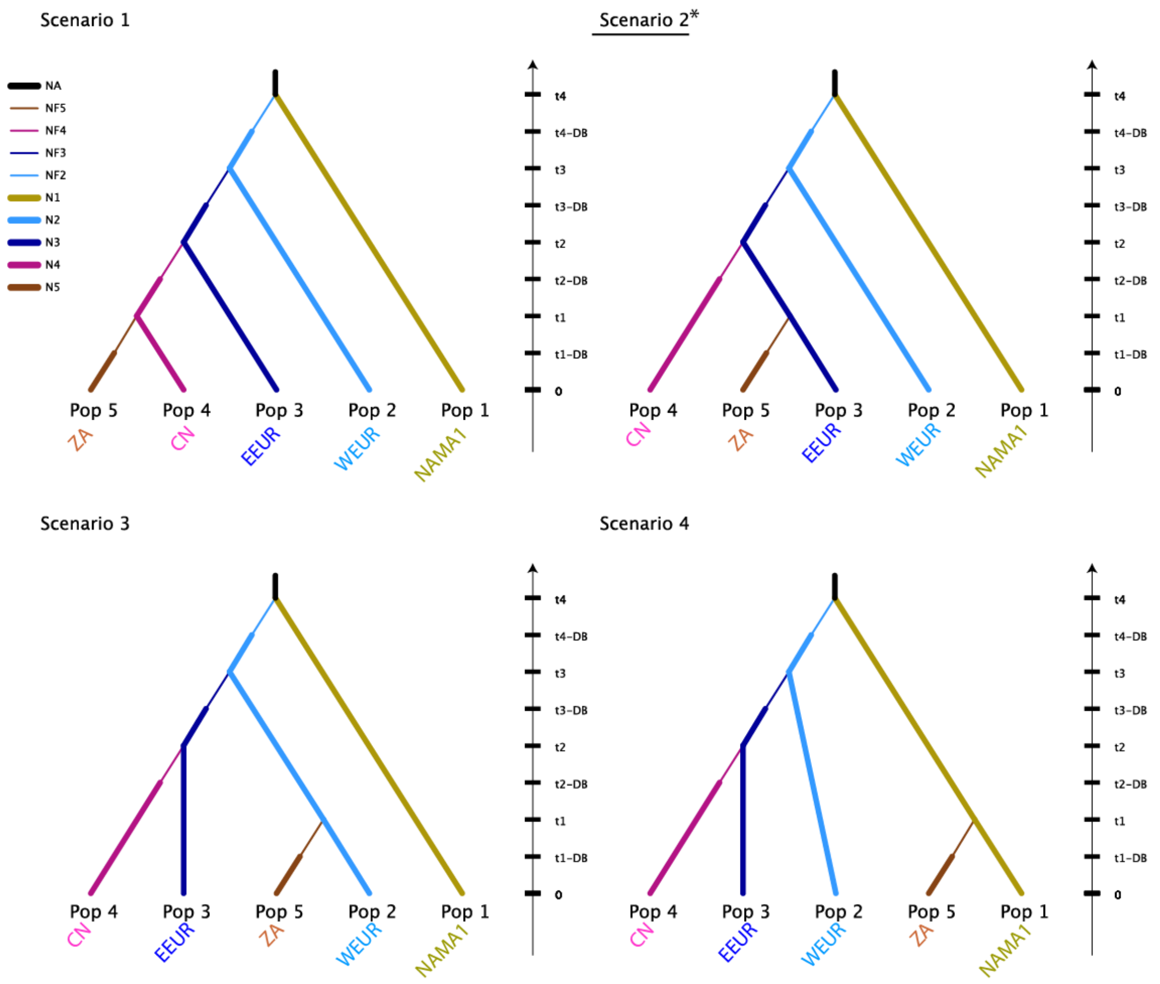


**Fig. S5.** Schematic representation of each scenario tested in the step 2 of the ABC analyses to decipher the worldwide invasion routes of *Plasmopara viticola forma specialis aestivalis*. It includes 4 possible scenarios describing the branching of the South African (ZA) cluster on each leaf of the best population tree identified in step 1 (SC11 in Fig. S5). For each analysis, the name of the most likely scenario is underlined and marked with an asterisk. Thin lines indicate bottlenecks. For parameter descriptions and priors see Table S12 and S13. Time is not to scale.

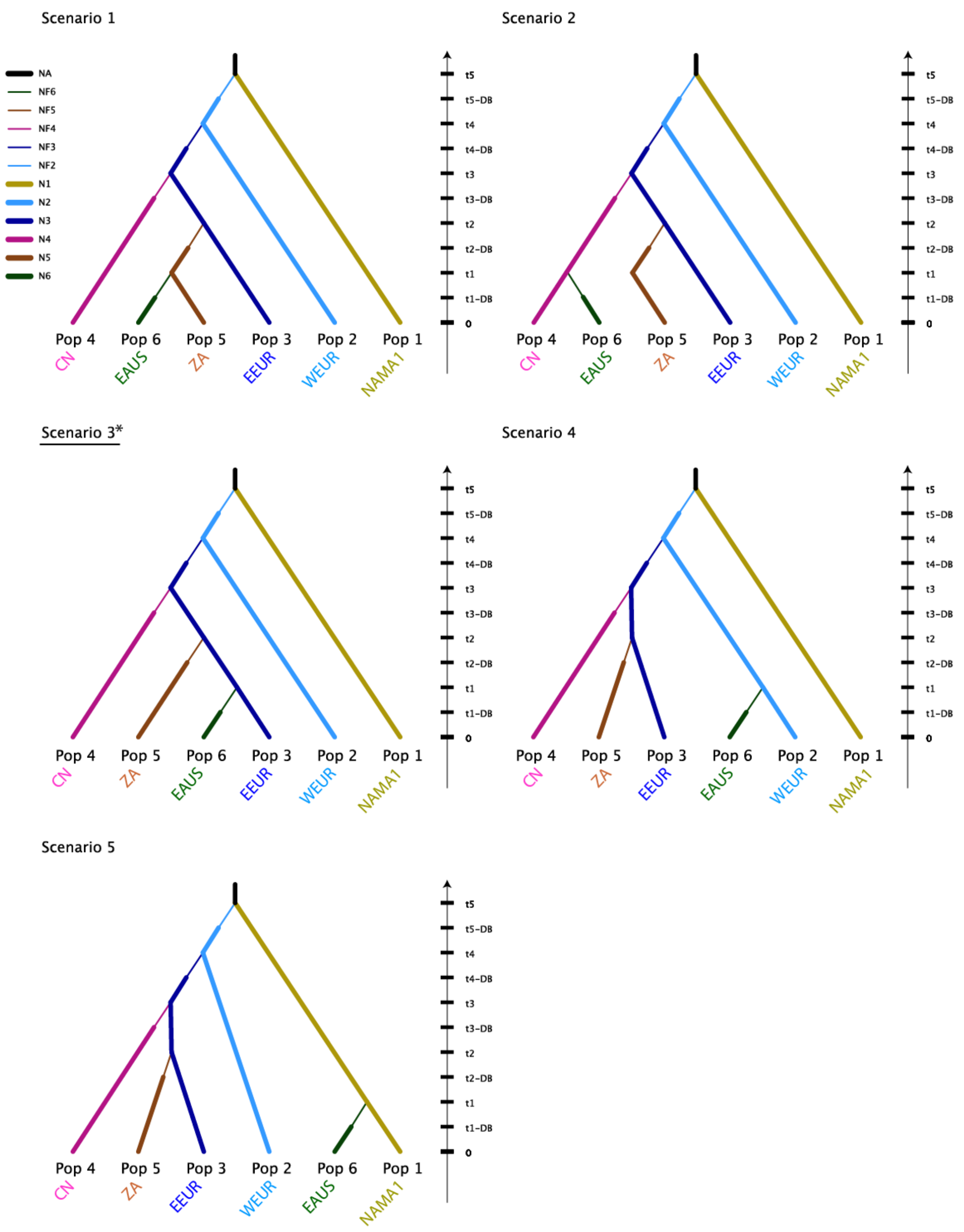


**Fig. S6.** Schematic representation of each scenario tested in the step 3 of the ABC analyses to decipher the worldwide invasion routes of *Plasmopara viticola forma specialis aestivalis*. It includes five possible scenarios describing the branching of the eastern Australian (EAUS) cluster on each leaf of the best population tree identified in step 2 (SC2 in Fig. S5). For each analysis, the name of the most likely scenario is underlined and marked with an asterisk. Thin lines indicate bottlenecks. For parameter descriptions and priors see Table S12 and S13. Time is not to scale.

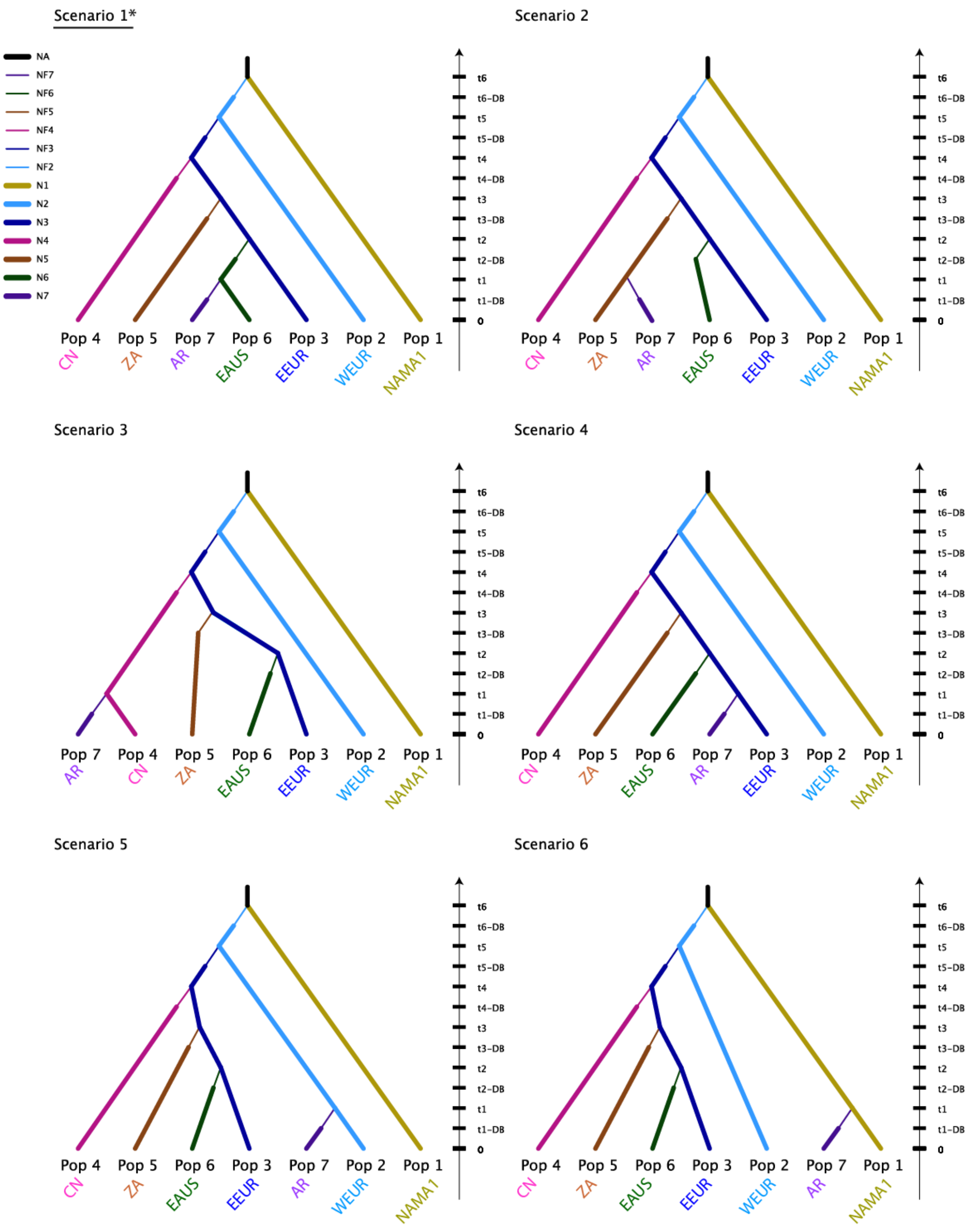


**Fig. S7.** Schematic representation of each scenario tested in the step 4 of the ABC analyses to decipher the worldwide invasion routes of *Plasmopara viticola forma specialis aestivalis*. It includes six possible scenarios describing the branching of the Argentan (AR) cluster on each leaf of the best population tree identified in step 3 (SC3 in Fig. S6). For each analysis, the name of the most likely scenario is underlined and marked with an asterisk. Thin lines indicate bottlenecks. For parameter descriptions and priors see Table S12 and S13. Time is not to scale.

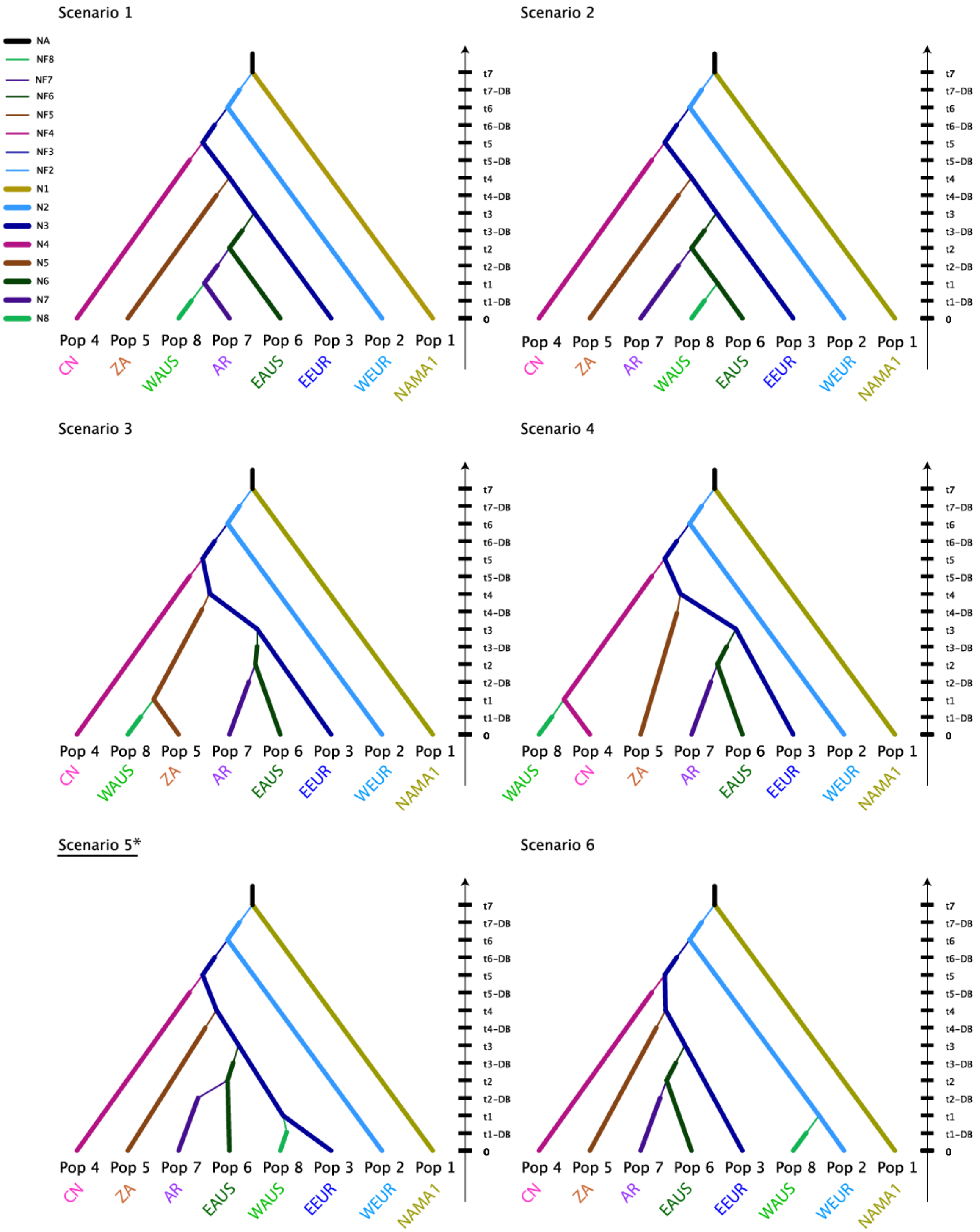


**Fig. S8** Schematic representation of each scenario tested in the step 5 of the ABC analyses to decipher the worldwide invasion routes of *Plasmopara viticola forma specialis aestivalis*. It includes seven possible scenarios describing the branching of the western Australian (WAUS) cluster on each leaf of the best population tree identified in step 4 (SC1 in Fig. S7). For each analysis, the name of the most likely scenario is underlined and marked with an asterisk. Thin lines indicate bottlenecks. For parameter descriptions and priors see Table S12 and S13. Time is not to scale. (1/2)

Scenario 7

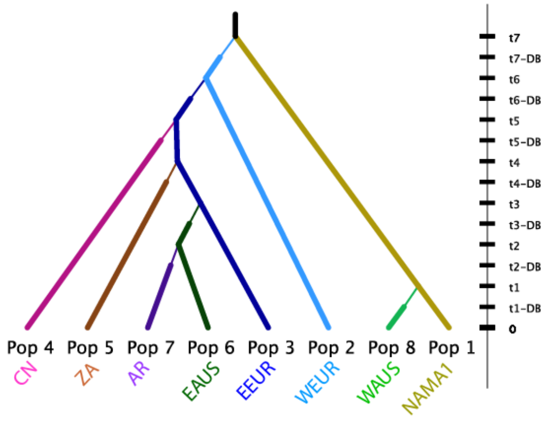


**Fig. S8.** Continued (2/2).

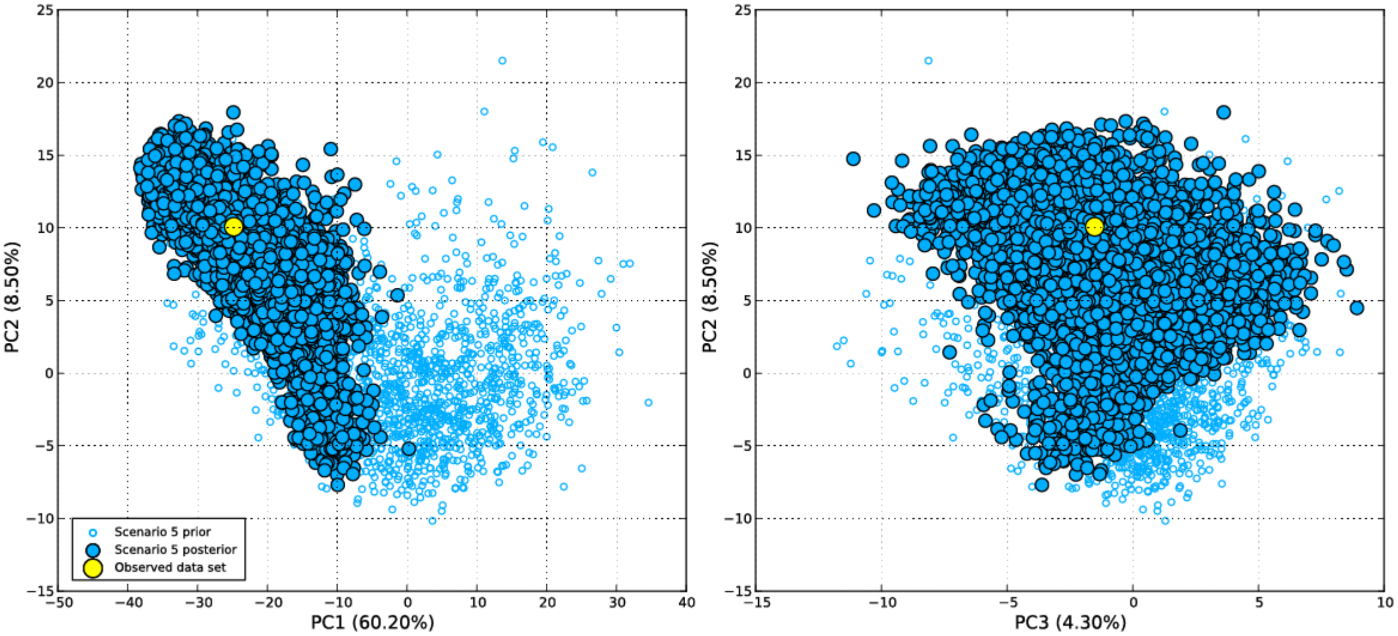


**Fig. S9.** Principal component analysis (PCA) conducted on the values obtained for 256 summary statistics used here as test quantities when processing ABC model-posterior checking for the final worldwide invasion scenario detailed in Fig. 4 and Fig. S8. The two panels display the scatter plots obtained for PC1 and 2 (left panel) and for PC2 and 3 (right panel) from the PCA accounting for up to 73% of the total variation. Small open blue circles show simulated datasets from the prior parameter distributions (10,000 simulations). Large filled blue circles show simulated datasets from posterior parameter distributions (10,000 simulations). The large yellow circle is the observed dataset. See the materials and methods text section for details.

**Table S1.** Genetic variation in different groups of *Plasmopara viticola* for each of the three following DNA fragments: β-tubulin (*tub,* n = 424, 499 bp), mitochondrial cytochrome-b (*cytb,* n = 1,299, 685 bp) and ribosomal 28S (r28S, n = 536, 703 bp). Values for each gene are separated by a slash (/); n*:* sample size; S: number of segregating sites; n_hap: number of haplotypes; H: haplotype diversity; π: per site nucleotide diversity; k: average number of differences among pairs of sequences.

| Formae speciales or population | | **n** | **S** | **n_hap** | **H** | **π (x 1e-3)** | **k** |
|---|---|---|---|---|---|---|---|
| *P. v. f. sp. vulpina* | Native | 8 / 4 / 4 | 3 / 0 / 0 | 3 / 1 / 1 | 0.71 / 0.00 / 0.00 | 3.15 / 0.00 / 0.00 | 1.57 / 0.00 / 0.00 |
| *P. v. f. sp. vinifera* | Native | 98 / 75 / 136 | 28 / 10 / 1 | 14 / 6 / 2 | 0.89 / 0.67 / 0.32 | 9.18 / 2.80 / 0.46 | 4.58 / 1.92 / 0.32 |
| *P. v. f. sp. quinquefolia* | Native | 6 / 1 / 6 | 1 / - / 3 | 2 / 1 / 3 | 0.53 / – / 0.73 | 1.07 / - / 1.99 | 0.53 / – / 1.40 |
| *P. v. f. sp. riparia* | Native | 112 / 24 / 162 | 13 / 4 / 0 | 12 / 4 / 1 | 0.44 / 0.60 / 0.00 | 2.83 / 1.76 / 0.00 | 1.41 / 1.20 / 0.00 |
| *P. v. f. sp. aestivalis* | | | | | | | |
| All populations | – | 624 / 1195 / 764 | 28 / 26 / 1 | 25 / 20 / 2 | 0.71 / 0.56 / 0.07 | 8.98 / 2.73 / 0.09 | 4.48 / 1.85 / 0.07 |
| North America | Native | 172 / 85 / 286 | 26 / 22 / 1 | 20 / 16 / 2 | 0.80 / 0.84 / 0.17 | 12.19 / 8.09 / 0.24 | 6.08 / 5.50 / 0.17 |
| Europe | Introduced | 258 / 995 / 260 | 6 / 4 / 0 | 5 / 4 / 1 | 0.56 / 0.48 / 0.00 | 3.24 / 1.48 / 0.00 | 1.62 / 1.02 / 0.00 |
| South Africa | Introduced | 72 / 47 / 90 | 2 / 4 / 0 | 3 / 3 / 1 | 0.39 / 0.45 / 0.00 | 1.46 / 1.47 / 0.00 | 0.73 / 1.01 / 0.00 |
| Australia | Introduced | 64 / 28 / 54 | 5 / 3 / 0 | 3 / 2 / 1 | 0.23 / 0.30 / 0.00 | 1.40 / 1.33 / 0.00 | 0.70 / 0.91 / 0.00 |
| China | Introduced | 36 / 30 / 50 | 2 / 3 / 0 | 2 / 2 / 1 | 0.36 / 0.29 / 0.00 | 1.43 / 1.26 / 0.00 | 0.71 / 0.86 / 0.00 |
| Argentina | Introduced | 22 / 10 / 24 | 2 / 3 / 0 | 2 / 2 / 1 | 0.17 / 0.20 / 0.00 | 0.69 / 0.88 / 0.00 | 0.35 / 0.60 / 0.00 |

**Table S2**: Genetic diversity for *Plasmopara viticola forma specialis aestivalis* populations based on eight microsatellite markers. n: sample size; $A_r$: allelic richness standardized for a minimum sample size of 14; $P_{Ar}$: private allelic richness standardized for a minimum sample size of 14; $H_o$: observed heterozygosity; $H_e$: unbiased expected heterozygosity (Nei, 1973); $F_{IS}$: fixation index (*: $P$ < 0.05; ** < 0.01; *** < 0.001). †Population in the native range closest to the introduced cultivated populations of the rest of the world vineyards.

| Continent | Group | Host | n | $A_r$ | $P_{Ar}$ | $H_e$ | $H_o$ | $F_{IS}$ |
|---|---|---|---|---|---|---|---|---|
| **North America** (Native) | NA1 | *V. labrusca* | 27 | 2.835 ± 1.222 | 0.581 ± 0.670 | 0.405 ± 0.289 | 0.187 ± 0.136 | 0.546*** |
| | NA2† | *V. aestivalis* | 21 | 3.676 ± 1.493 | 0.600 ± 0.987 | 0.589 ± 0.204 | 0.399 ± 0.238 | 0.329*** |
| | NA3 | *V. vinifera* | 16 | 2.167 ± 1.011 | 0.273 ± 0.411 | 0.310 ± 0.266 | 0.306 ± 0.279 | 0.016 |
| | NA4 | *V. vinifera* | 111 | 3.607 ± 1.036 | 0.516 ± 0.398 | 0.568 ± 0.195 | 0.386 ± 0.169 | 0.322*** |
| **Europe** (Introduced) | WEU | *V. vinifera* | 715 | 2.316 ± 0.633 | 0.035 ± 0.049 | 0.3910 ± 0.199 | 0.356 ± 0.218 | 0.09*** |
| | EEU | - | 265 | 2.391 ± 0.630 | 0.030 ± 0.032 | 0.356 ± 0.184 | 0.318 ± 0.224 | 0.108*** |
| **South Africa** (Introduced) | ZA | - | 67 | 2.250 ± 0.598 | 0.078 ± 0.186 | 0.381 ± 0.219 | 0.339 ± 0.279 | 0.11 |
| **Australia** (Introduced) | WAU | - | 14 | 1.727 ± 1.142 | 0.000 ± 0.000 | 0.203 ± 0.286 | 0.232 ± 0.323 | -0.148 |
| | EAU | - | 36 | 2.333 ± 0.753 | 0.001 ± 0.002 | 0.411 ± 0.182 | 0.376 ± 0.235 | 0.09 |
| **China** (Introduced) | CN | - | 25 | 2.042 ± 0.428 | 0.012 ± 0.035 | 0.324 ± 0.178 | 0.366 ± 0.217 | -0.131 |
| **Argentina** (Introduced) | AR | - | 17 | 1.679 ± 0.457 | 0.000 ± 0.000 | 0.285 ± 0.230 | 0.347 ± 0.373 | -0.222 |

**Table S3:** Genetic diversity in *Plasmopara viticola forma specialis aestivalis* populations based on the 32 microsatellite marker dataset. n: clone-corrected sample size; $A_r$: allelic richness standardized for a minimum sample size of two; $P_{Ar}$: private allelic richness standardized for a minimum sample size of two; $H_o$: observed heterozygosity; $H_e$: unbiased expected heterozygosity (Nei, 1973); $F_{IS}$: fixation index (*: $P < 0.05$; ** < 0.01; *** < 0.001). Population acronyms are as follows: North American *P. viticola* f. sp. *aestivalis* strains collected on wild *Vitis labrusca* (NA1), *V. aestivalis* group 1 (NA2) and group 2 (NA3), and North American strains collected on cultivated *V. vinifera* (NA4). Strains in the rest of the world were collected on cultivated *V. vinifera* including Western and Eastern Europe (WEU and WEU), South Africa (ZA), Western and Eastern Australia (WAU and EAU), China (CN), and Argentina (AR). †Population in the native range closest to the introduced cultivated populations of the rest of the world vineyards.

| Continent | Group | Range | Host | n | $A_r \pm SD$ | $P_{Ar} \pm SD$ | $H_o$ | $H_e$ | $F_{IS}$ |
|---|---|---|---|---|---|---|---|---|---|
| North America | NA1 | Native | *V. labrusca* | 15 | 1.23 ± 0.23 | 0.30 ± 0.40 | 0.30 ± 0.31 | 0.30 ± 0.23 | -0.034 |
| - | NA2† | Native | *V. aestivalis* | 5 | 1.49 ± 0.28 | 0.26 ± 0.34 | 0.40 ± 0.34 | 0.47 ± 0.30 | 0.156 |
| - | NA3 | Native | *V. aestivalis* | 4 | 1.30 ± 0.26 | 0.16 ± 0.27 | 0.46 ± 0.46 | 0.32 ± 0.26 | -0.600*** |
| - | NA4 | Native | *V. vinifera* | 27 | 1.38 ± 0.19 | 0.19 ± 0.25 | 0.33 ± 0.24 | 0.39 ± 0.21 | 0.161*** |
| Europe | WEU | Introduced | *V. vinifera* | 48 | 1.35 ± 0.23 | 0.03 ± 0.05 | 0.30 ± 0.23 | 0.34 ± 0.23 | 0.121*** |
| - | EEU | Introduced | *V. vinifera* | 19 | 1.37 ± 0.24 | 0.03 ± 0.06 | 0.36 ± 0.27 | 0.36 ± 0.24 | 0.004 |
| South Africa | ZA | Introduced | *V. vinifera* | 12 | 1.32 ± 0.25 | 0.04 ± 0.10 | 0.40 ± 0.39 | 0.32 ± 0.25 | -0.279*** |
| Australia | WAU | Introduced | *V. vinifera* | 7 | 1.16 ± 0.25 | 0.01 ± 0.01 | 0.25 ± 0.43 | 0.15 ± 0.25 | -0.833*** |
| - | EAU | Introduced | *V. vinifera* | 5 | 1.25 ± 0.25 | 0.03 ± 0.10 | 0.31 ± 0.42 | 0.24± 0.26 | -0.318 |
| China | CN | Introduced | *V. vinifera* | 24 | 1.29 ± 0.22 | 0.02 ± 0.05 | 0.36 ± 0.31 | 0.28 ± 0.22 | -0.278*** |
| Argentina | AR | Introduced | *V. vinifera* | 8 | 1.22 ± 0.22 | 0.02 ± 0.04 | 0.25 ± 0.31 | 0.23 ± 0.22 | -0.100 |

**Table S4**: Pairwise differentiation estimated as $F_{ST}$ among *Plasmopara viticola* populations using the eight-microsatellite-marker dataset (dataset 1). The significance of the population differentiation was tested using a permutation test (100 permutations). The upper-triangular and lower-triangular matrices show the $F_{ST}$ values and their significance level, respectively (***: P-value < 0.001). Population acronyms are as follows: North American *P. viticola* strains collected on wild *Vitis labrusca* (NA1), North American strains collected on *V. aestivalis* (NA2), North American strains collected on cultivated *V. vinifera* group 1 (NA3) and group 2 (NA4). Strains in the rest of the world were collected on cultivated *V. vinifera* including Western and Eastern Europe (WEU and WEU), South Africa (ZA), Western and Eastern Australia (WAU and EAU), China (CN), and Argentina (AR).

| Microsatellite dataset 1 (8 loci) | | | | | | | | | | | |
|---|---|---|---|---|---|---|---|---|---|---|---|
| **Group** | **NA1** | **NA2** | **NA3** | **NA4** | **W-EU** | **E-EU** | **ZA** | **W-AU** | **E-AU** | **CN** | **AR** |
| **NA1** | | 0.25 | 0.50 | 0.23 | 0.45 | 0.50 | 0.48 | 0.57 | 0.42 | 0.53 | 0.57 |
| **NA2** | *** | | 0.33 | 0.12 | 0.29 | 0.33 | 0.26 | 0.32 | 0.25 | 0.31 | 0.31 |
| **NA3** | *** | *** | | 0.36 | 0.31 | 0.34 | 0.36 | 0.64 | 0.39 | 0.37 | 0.48 |
| **NA4** | *** | *** | *** | | 0.38 | 0.39 | 0.33 | 0.36 | 0.30 | 0.35 | 0.38 |
| **W-EU** | *** | *** | *** | *** | | 0.02 | 0.08 | 0.30 | 0.12 | 0.06 | 0.14 |
| **E-EU** | *** | *** | *** | *** | *** | | 0.10 | 0.36 | 0.19 | 0.03 | 0.17 |
| **ZA** | *** | *** | *** | *** | *** | *** | | 0.24 | 0.10 | 0.15 | 0.17 |
| **W-AU** | *** | *** | *** | *** | *** | *** | *** | | 0.26 | 0.46 | 0.44 |
| **E-AU** | *** | *** | *** | *** | *** | *** | *** | *** | | 0.24 | 0.20 |
| **CN** | *** | *** | *** | *** | *** | *** | *** | *** | *** | | 0.20 |
| **AR** | *** | *** | *** | *** | *** | *** | *** | *** | *** | *** | |

**Table S5**: Pairwise differentiation estimated as $F_{ST}$ among *Plasmopara viticola* populations using the microsatellite 32-microsatellite-marker dataset (dataset 2). The significance of the population differentiation was tested using a permutation test (100 permutations). The upper-triangular and lower-triangular matrices show the $F_{ST}$ values and their significance level, respectively (***: P-value < 0.001). Bold values show the native vs. introduced pairwise comparisons. Population acronyms are as follows: North American *P. viticola* strains collected on wild *Vitis labrusca* (NA1), *V. aestivalis* group 1 (NA2) and group 2 (NA3), and North American strains collected on cultivated *V. vinifera* (NA4). Strains in the rest of the world were collected on cultivated *V. vinifera* including Western and Eastern Europe (WEU and WEU), South Africa (ZA), Western and Eastern Australia (WAU and EAU), China (CN), and Argentina (AR).

| Microsatellite dataset 2 (32 loci) | | | | | | | | | | | |
|---|---|---|---|---|---|---|---|---|---|---|---|
| **Group** | **NA1** | **NA2** | **NA3** | **NA4** | **W-EU** | **E-EU** | **ZA** | **W-AU** | **E-AU** | **CN** | **AR** |
| **NA1** | | 0.47 | 0.55 | 0.40 | 0.53 | 0.55 | 0.58 | **0.67** | 0.62 | 0.62 | 0.66 |
| **NA2** | *** | | 0.33 | 0.28 | 0.38 | 0.33 | 0.41 | 0.49 | 0.44 | 0.44 | 0.46 |
| **NA3** | *** | *** | | 0.24 | 0.49 | 0.49 | 0.54 | **0.71** | 0.62 | 0.59 | 0.60 |
| **NA4** | *** | *** | *** | | 0.41 | 0.41 | 0.43 | 0.52 | 0.45 | 0.48 | 0.47 |
| **W-EU** | *** | *** | *** | *** | | **0.07** | 0.11 | 0.30 | 0.24 | 0.17 | 0.19 |
| **E-EU** | *** | *** | *** | *** | *** | | **0.08** | 0.31 | 0.21 | 0.10 | 0.14 |
| **ZA** | *** | *** | *** | *** | *** | *** | | 0.36 | 0.29 | 0.21 | 0.24 |
| **W-AU** | *** | *** | *** | *** | *** | *** | *** | | **0.50** | 0.40 | **0.55** |
| **E-AU** | *** | *** | *** | *** | *** | *** | *** | *** | | 0.40 | 0.31 |
| **CN** | *** | *** | *** | *** | *** | *** | *** | *** | *** | | 0.33 |
| **AR** | *** | *** | *** | *** | *** | *** | *** | *** | *** | *** | |

**Table S6**: Test of pairwise differences in allelic richness ($A_r$) between *Plasmopara viticola* populations, using the microsatellite dataset 1, based on Wilcoxon signed rank tests. The upper-triangular and lower-triangular matrices show the *P-values* and their significance, respectively (*: *P* < 0.05; ** < 0.01; *** < 0.001; ns: non significant). Bold values show the native vs. introduced pairwise comparisons. Population acronyms are as follows: North American *P. viticola* strains collected on wild *Vitis labrusca* (NA1), North American strains collected on *V. aestivalis* (NA2), North American strains collected on cultivated *V. vinifera* group 1 (NA3) and group 2 (NA4). Strains in the rest of the world were collected on cultivated *V. vinifera* including Western and Eastern Europe (WEU and WEU), South Africa (ZA), Western and Eastern Australia (WAU and EAU), China (CN), and Argentina (AR). †Population in the native range closest to the introduced cultivated populations of the rest of the world vineyards.

| **Microsatellite dataset 1 (8 markers)** | | | | | | | | | | | |
|---|---|---|---|---|---|---|---|---|---|---|---|
| **Group** | NA1 | **NA2†** | NA3 | NA4 | WEU | EEU | ZA | WAU | EAU | CN | AR |
| NA1 | | **0.078** | 0.313 | 0.016 | 0.461 | 0.547 | 0.195 | 0.148 | 0.383 | 0.195 | 0.039 |
| **NA2†** | ns | | **0.008** | 0.844 | **0.008** | **0.008** | **0.008** | **0.016** | **0.008** | **0.008** | **0.008** |
| NA3 | ns | ** | | 0.008 | 0.461 | 0.641 | 0.844 | 0.205 | 0.461 | 0.547 | 0.208 |
| NA4 | * | **ns** | ** | | 0.016 | 0.016 | 0.016 | 0.016 | 0.023 | 0.016 | 0.008 |
| WEU | ns | ** | ns | * | | 0.641 | 1.000 | 0.078 | 0.844 | 0.109 | 0.016 |
| EEU | ns | ** | ns | * | ns | | 0.641 | 0.109 | 0.945 | 0.023 | 0.039 |
| ZA | ns | ** | ns | * | ns | ns | | 0.250 | 0.945 | 0.461 | 0.016 |
| WAU | ns | * | ns | * | ns | ns | ns | | 0.042 | 0.250 | 0.800 |
| EAU | ns | ** | ns | * | ns | ns | ns | * | | 0.076 | 0.042 |
| CN | ns | ** | ns | * | ns | * | ns | ns | ns | | 0.109 |
| AR | * | ** | ns | ** | * | * | * | ns | * | ns | |

**Table S7**: Test of pairwise differences in allelic richness ($A_r$) between *Plasmopara viticola* populations, using the microsatellite dataset 2, based on Wilcoxon signed rank tests. The upper-triangular and lower-triangular matrices show the *P-values* and their significance, respectively (*: $P < 0.05$; ** < 0.01; *** < 0.001; ns: non significant). Bold values show the native vs. introduced pairwise comparisons. Population acronyms are as follows: North American *P. viticola* strains collected on wild *Vitis labrusca* (NA1), *V. aestivalis* group 1 (NA2) and group 2 (NA3), and North American strains collected on cultivated *V. vinifera* (NA4). Strains in the rest of the world were collected on cultivated *V. vinifera* including Western and Eastern Europe (WEU and WEU), South Africa (ZA), Western and Eastern Australia (WAU and EAU), China (CN), and Argentina (AR). †Population in the native range closest to the introduced cultivated populations of the rest of the world vineyards.

| **Microsatellite dataset 2 (32 markers)** | | | | | | | | | | | |
|---|---|---|---|---|---|---|---|---|---|---|---|
| **Group** | NA1 | **NA2†** | NA3 | NA4 | WEU | EEU | ZA | WAU | EAU | CN | AR |
| NA1 | | 0.003 | 1.000 | 0.078 | **0.473** | **0.279** | **0.728** | **0.028** | **0.464** | **0.741** | **0.244** |
| **NA2†** | ** | | 0.006 | 0.019 | **0.039** | **0.042** | **0.018** | **0.000** | **0.000** | **0.002** | **0.001** |
| NA3 | ns | ** | | 0.116 | **0.494** | **0.215** | **0.721** | **0.033** | **0.282** | **0.936** | **0.247** |
| NA4 | ns | * | ns | | **0.495** | **0.717** | **0.315** | **0.002** | **0.011** | **0.030** | **0.003** |
| WEU | **ns** | * | **ns** | **ns** | | 0.374 | 0.432 | 0.007 | 0.067 | 0.343 | 0.014 |
| EEU | **ns** | * | **ns** | **ns** | ns | | 0.083 | 0.002 | 0.029 | 0.028 | 0.001 |
| ZA | **ns** | * | **ns** | **ns** | ns | ns | | 0.027 | 0.218 | 0.597 | 0.112 |
| W-AU | **ns** | *** | * | ** | ** | ** | * | | 0.074 | 0.030 | 0.295 |
| E-AU | **ns** | *** | **ns** | * | ns | * | ns | ns | | 0.362 | 0.745 |
| CN | **ns** | ** | **ns** | * | ns | * | ns | * | ns | | 0.127 |
| AR | **ns** | ** | **ns** | ** | * | ** | ns | ns | ns | ns | |

**Table S8**: Test of pairwise differences in private allelic richness ($P_{Ar}$) between *Plasmopara viticola* populations, using the microsatellite dataset 1 (eight markers), based on Wilcoxon signed rank tests. The upper-triangular and lower-triangular matrices show the *P-values* and their significance levels, respectively (*: $P < 0.05$; ** < 0.01; *** < 0.001; ns: non-significant). Bold values show the native vs. introduced pairwise comparisons. †Population acronyms are as follows: North American *P. viticola* strains collected on wild *Vitis labrusca* (NA1), North American strains collected on *V. aestivalis* (NA2), North American strains collected on cultivated *V. vinifera* group 1 (NA3) and group 2 (NA4). Strains in the rest of the world were collected on cultivated *V. vinifera* including Western and Eastern Europe (WEU and WEU), South Africa (ZA), Western and Eastern Australia (WAU and EAU), China (CN), and Argentina (AR). †Population in the native range closest to the introduced cultivated populations of the rest of the world vineyards.

| Microsatellite dataset 1 (eight markers) | | | | | | | | | | | |
|---|---|---|---|---|---|---|---|---|---|---|---|
| **Group** | NA1 | **NA2†** | NA3 | NA4 | W-EU | E-EU | ZA | W-AU | E-AU | CN | AR |
| NA1 | | 0.834 | 0.208 | 0.945 | 0.108 | 0.076 | 0.076 | 0.059 | 0.059 | 0.059 | 0.059 |
| **NA2†** | ns | | 0.447 | 0.844 | 0.108 | 0.195 | 0.109 | 0.036 | 0.036 | 0.076 | 0.036 |
| NA3 | ns | ns | | 0.461 | 0.554 | 0.401 | 0.402 | 0.181 | 0.181 | 0.201 | 0.181 |
| NA4 | ns | ns | ns | | 0.008 | 0.016 | 0.023 | 0.008 | 0.008 | 0.008 | 0.008 |
| W-EU | ns | ns | ns | ** | | 0.933 | 0.554 | 0.022 | 0.022 | 0.022 | 0.022 |
| E-EU | ns | ns | ns | * | ns | | 0.529 | 0.036 | 0.036 | 0.142 | 0.036 |
| ZA | ns | ns | ns | * | ns | ns | | 0.036 | 0.115 | 0.293 | 0.036 |
| W-AU | ns | * | ns | ** | * | * | * | | 1.000 | 1.000 | 1.000 |
| E-AU | ns | * | ns | ** | * | * | ns | ns | | 1.000 | 1.000 |
| CN | ns | ns | ns | ** | * | ns | ns | ns | ns | | 1.000 |
| AR | ns | * | ns | ** | * | * | * | ns | ns | ns | |

**Table S9**: Test of pairwise differences in private allelic richness ($P_{Ar}$) between *Plasmopara viticola* populations, using the microsatellite dataset 2 (32 markers), based on Wilcoxon signed rank tests. The upper-triangular and lower-triangular matrices show the *P-values* and their significance levels, respectively (*: *P* < 0.05; ** < 0.01; *** < 0.001; ns: non-significant). Bold values show the native vs. introduced pairwise comparisons. Population acronyms are as follows: North American *P. viticola* strains collected on wild *Vitis labrusca* (NA1), *V. aestivalis* group 1 (NA2) and group 2 (NA3), and North American strains collected on cultivated *V. vinifera* (NA4). Strains in the rest of the world were collected on cultivated *V. vinifera* including Western and Eastern Europe (WEU and WEU), South Africa (ZA), Western and Eastern Australia (WAU and EAU), China (CN), and Argentina (AR). †Population in the native range closest to the introduced cultivated populations of the rest of the world vineyards.

| Microsatellite dataset 2 (32 markers) | | | | | | | | | | | |
|---|---|---|---|---|---|---|---|---|---|---|---|
| **Group** | NA1 | **NA2†** | NA3 | NA4 | W-EU | E-EU | ZA | W-AU | E-AU | CN | AR |
| NA1 | | 0.843 | 0.080 | 0.099 | **0.000** | **0.001** | **0.001** | **0.000** | **0.000** | **0.001** | **0.001** |
| **NA2†** | ns | | 0.119 | 0.416 | **0.001** | **0.002** | **0.003** | **0.000** | **0.000** | **0.001** | **0.001** |
| NA3 | ns | ns | | 0.493 | **0.019** | **0.046** | **0.018** | **0.001** | **0.011** | **0.032** | **0.008** |
| NA4 | ns | ns | ns | | **0.000** | **0.001** | **0.005** | **0.000** | **0.001** | **0.000** | **0.001** |
| W-EU | *** | ** | * | *** | | 1.000 | 0.925 | 0.010 | 0.052 | 0.679 | 0.218 |
| E-EU | ** | ** | * | ** | ns | | 0.906 | 0.018 | 0.177 | 0.421 | 0.127 |
| ZA | ** | ** | * | ** | ns | ns | | 0.055 | 0.443 | 0.754 | 0.224 |
| W-AU | *** | *** | *** | *** | ** | * | ns | | 0.343 | 0.006 | 0.476 |
| E-AU | *** | *** | * | *** | ns | ns | ns | ns | | 0.066 | 0.649 |
| CN | ** | *** | * | *** | ns | ns | ns | ** | ns | | 0.286 |
| AR | ** | *** | ** | *** | ns | ns | ns | ns | ns | ns | |

**Table S10**: Test of pairwise differences in expected heterozygosity ($H_E$) between *Plasmopara viticola* populations, using the microsatellite dataset 1 (eight markers), based on Wilcoxon signed rank tests. The upper-triangular and lower-triangular matrices show the *P-values* and the significance of the corresponding *P-values,* respectively (*: $P$ < 0.05; ** < 0.01; *** < 0.001; ns: non-significant). Bold values show the native vs. introduced pairwise comparisons. Population acronyms are as follows: North American *P. viticola* strains collected on wild *Vitis labrusca* (NA1), North American strains collected on *V. aestivalis* (NA2), North American strains collected on cultivated *V. vinifera* group 1 (NA3) and group 2 (NA4). Strains in the rest of the world were collected on cultivated *V. vinifera* including Western and Eastern Europe (WEU and WEU), South Africa (ZA), Western and Eastern Australia (WAU and EAU), China (CN), and Argentina (AR). †Population in the native range closest to the introduced cultivated populations of the rest of the world vineyards.

| Microsatellite dataset 1 (eight markers) | | | | | | | | | | | |
|---|---|---|---|---|---|---|---|---|---|---|---|
| **Group** | NA1 | **NA2†** | NA3 | NA4 | WEU | EEU | ZA | WAU | EAU | CN | AR |
| NA1 | | 0.148 | 0.461 | 0.148 | 0.945 | 0.641 | 0.844 | 0.400 | 1.000 | 0.641 | 0.250 |
| **NA2†** | ns | | 0.055 | 0.742 | 0.039 | 0.023 | 0.109 | 0.008 | 0.055 | 0.023 | 0.008 |
| NA3 | ns | ns | | 0.023 | 0.250 | 0.547 | 0.641 | 0.205 | 0.195 | 0.945 | 0.933 |
| NA4 | ns | ns | * | | 0.109 | 0.055 | 0.148 | 0.016 | 0.148 | 0.078 | 0.008 |
| WEU | ns | * | ns | ns | | 0.383 | 1.000 | 0.078 | 0.641 | 0.383 | 0.195 |
| EEU | ns | * | ns | ns | ns | | 0.844 | 0.078 | 0.313 | 0.547 | 0.383 |
| ZA | ns | ns | ns | ns | ns | ns | | 0.250 | 0.742 | 0.547 | 0.195 |
| WAU | ns | ** | ns | * | ns | ns | ns | | 0.055 | 0.313 | 0.447 |
| EAU | ns | ns | ns | ns | ns | ns | ns | ns | | 0.195 | 0.148 |
| CN | ns | * | ns | ns | ns | ns | ns | ns | ns | | 0.547 |
| AR | ns | ** | ns | ** | ns | ns | ns | ns | ns | ns | |

**Table S11**: Test of pairwise differences in expected heterozygosity ($H_E$) between *Plasmopara viticola* populations, using the microsatellite dataset 2 (32 markers), based on Wilcoxon signed rank tests. The upper-triangular and lower-triangular matrices show the *P-values* and the significance of the corresponding *P-values,* respectively (*: $P < 0.05$; ** < 0.01; *** < 0.001; ns: non-significant). Bold values show the native vs. introduced pairwise comparisons. Population acronyms are as follows: North American *P. viticola* strains collected on wild *Vitis labrusca* (NA1), *V. aestivalis* group 1 (NA2) and group 2 (NA3), and North American strains collected on cultivated *V. vinifera* (NA4). Strains in the rest of the world were collected on cultivated *V. vinifera* including Western and Eastern Europe (WEU and WEU), South Africa (ZA), Western and Eastern Australia (WAU and EAU), China (CN), and Argentina (AR). †Population in the native range closest to the introduced cultivated populations of the rest of the world vineyards.

| Microsatellite dataset 2 (32 markers) | | | | | | | | | | | |
|---|---|---|---|---|---|---|---|---|---|---|---|
| **Group** | NA1 | **NA2†** | NA3 | NA4 | WEU | EEU | ZA | WAU | EAU | CN | AR |
| NA1 | | 0.008 | 0.829 | 0.033 | **0.476** | **0.289** | **0.707** | **0.018** | **0.516** | **0.721** | **0.210** |
| **NA2†** | ** | | 0.011 | 0.077 | **0.058** | **0.063** | **0.025** | **0.000** | **0.001** | **0.003** | **0.001** |
| NA3 | ns | * | | 0.176 | **0.779** | **0.434** | **0.960** | **0.019** | **0.139** | **0.540** | **0.162** |
| NA4 | * | ns | ns | | **0.432** | **0.612** | **0.266** | **0.001** | **0.006** | **0.016** | **0.004** |
| WEU | **ns** | **ns** | **ns** | **ns** | | 0.304 | 0.589 | 0.003 | 0.049 | 0.243 | 0.017 |
| EEU | **ns** | **ns** | **ns** | **ns** | ns | | 0.110 | 0.001 | 0.021 | 0.015 | 0.002 |
| ZA | **ns** | * | **ns** | **ns** | ns | ns | | 0.012 | 0.162 | 0.402 | 0.095 |
| WAU | * | *** | * | *** | ** | *** | * | | 0.042 | 0.015 | 0.173 |
| EAU | **ns** | *** | **ns** | ** | * | * | ns | * | | 0.360 | 0.875 |
| CN | **ns** | ** | **ns** | * | ns | * | ns | * | ns | | 0.184 |
| AR | **ns** | ** | **ns** | ** | * | ** | ns | ns | ns | ns | |

**Table S12. Description of the competing scenarios and results of the five successive steps of random forest classification analysis (ABC-RF) to infer the invasion history of *Plasmopara viticola* based on the genetic variation at 32 microsatellite markers.** For each ABC analysis, an ABC-RF analysis was run 10 times using a training set ranging from 10 to 50 thousand simulations and a forest of 500 or 1,000 trees. The size of the training set and number of trees used in each step was increased until the 10 ABC-RF replicates all converged. The best (most likely) scenario identified at each step is shown in bold and marked with a (*) symbol. Alternative scenario(s) receiving significant support (≥15% of the votes) are indicated with the symbol (†). Prior error rates, proportion of votes, and posterior probability values are averaged over the 10 replicates. Stepwise introduction is indicated as [source] > [derived] populations.

| Scenarios tested in 5 steps | Training set | Decision trees | Prior error rate (%) | RF votes (%) | Post. Prob ± SD |
|---|---|---|---|---|---|
| **Step 1** (18 scenarios, 70 summary statistics):<br>North America *aestivalis 1 (NA2) to WEUR / EEUR / CN* | 50,000 | 1,000 | 38% | | |
| S1: Independent introductions from NA2 to CN, then WEUR, then EEUR | | | | 0.1% | – |
| S2: Independent introductions from NA2 to CN, then EEUR, then WEUR | | | | 0.1% | – |
| S3: Independent introductions from NA2 to EEUR, then WEUR, then CN | | | | 0.2% | – |
| S4: Independent introductions from NA2 to WEUR, then EEUR, then CN | | | | 0.1% | – |
| S5: Introduction from NA2 > WEUR > EEUR, CN admixed btw. WEUR-EEUR | | | | 8.6% | – |
| S6: Introduction from NA2 > EEUR > WEUR, CN admixed btw. WEUR-EEUR | | | | 3.9% | – |
| S7†: Introduction from NA2 > CN > WEUR, EEUR admixed btw. CN-WEUR | | | | 16.0% | – |
| S8†: Introduction from NA2 > WEUR > CN, EEUR admixed btw. CN-WEUR | | | | 17.5% | – |
| S9: Introduction from NA2 > EEUR > CN, WEUR admixed btw. CN-EEUR | | | | 5.0% | – |
| S10: Introduction from NA2 > CN > EEUR, WEUR admixed btw. CN-EEUR | | | | 6.2% | – |
| **S11*: Introduction from NA2 > WEUR > EEUR > CN** | | | | **20.1%** | **0.42 ± 0.01** |
| S12: Introduction from NA2 > EEUR > WEUR > CN | | | | 0.5% | – |
| S13: Introduction from NA2 > CN > WEUR > EEUR | | | | 2.9% | – |
| S14: Introduction from NA2 > WEUR > CN > EEUR | | | | 3.4% | – |
| S15: Introduction from NA2 > EEUR > CN > WEUR | | | | 0.4% | – |
| S16: Introduction from NA2 > CN > EEUR > WEUR | | | | 13.5% | – |
| S17: Independent introduction of WEUR, then CN; EEUR admixed btw. WEUR-CN | | | | 0.7% | – |
| S18: Independent introduction of CN, then WEUR; EEUR admixed btw. WEUR-CN | | | | 0.8% | – |
| **Step 2** (4 scenarios based on S11 of step 1, 100 summary statistics):<br>Introduction of South Africa (ZA) from | 10,000 | 500 | 9% | | |
| S1: CN | | | | 8% | – |
| **S2*: EEUR** | | | | **50%** | **0.78 ± 0.03** |
| S3†: WEUR | | | | 42% | – |
| S4: NA2 | | | | 0% | – |
| **Step 3** (5 scenarios based on S2 of step 2, 144 summary statistics):<br>Introduction of East Australia (EAUS) from … | 20,000 | 500 | 12% | | |
| S1: ZA | | | | 11% | – |
| S2: CN | | | | 11% | – |
| **S3*: EEUR** | | | | **39%** | **0.71 ± 0.02** |
| S4†: WEUR | | | | 29% | – |
| S5: NA2 | | | | 10% | – |
| **Step 4** (6 scenarios based on S3 of step 3, 195 summary statistics):<br>Introduction of Argentina (AR) from | 10,000 | 500 | 12% | | |
| **S1*: EAUS** | | | | **48%** | **0.70 ± 0.02** |
| S2: ZA | | | | 12% | – |
| S3: CN | | | | 5% | – |
| S4†: EEUR | | | | 23% | – |
| S5: WEUR | | | | 9% | – |
| S6: NA2 | | | | 2% | – |
| **Step 5** (7 scenarios based on S1 of step 4, 256 summary statistics):<br>Introduction of West Australia (WAUS) from | 10,000 | 1 000 | 7% | | |
| S1: AR | | | | 6% | – |
| S2†: EAUS | | | | 18% | – |
| S3†: ZA | | | | 17% | – |
| S4†: CN | | | | 18% | – |
| **S5*: EEUR** | | | | **23%** | **0.73 ± 0.02** |
| S6: WEUR | | | | 14% | – |
| S7: NA2 | | | | 4% | – |

**Table S13**. Prior distributions for the demographic and mutation model parameters used in ABC-RF analyses processed to retrace the worldwide invasion routes of *Plasmopara viticola.* Demographic parameters include the effective size ($N_i$) for the *i*-populations, where *i* ranges from 1 to 8 and include also the ancestral population NA; the splitting time ($T_{j\ or\ k}$, in generations, with *j* ranging from 1 to 7 and *k* ranging from 2 to 6), duration of the bottleneck (DB), and effective size during the bottleneck ($NF_i$). The parameters for the microsatellite mutation model include the mean and individual locus mutation rate ($\mu$), the parameter of the geometric distribution (*P*); and the single nucleotide insertion rate (*SNI*). Three types of prior distributions are used: uniform, UN~[Min–Max]; gamma, GA~[Min, Max, Shape]; and log-uniform, LU~[Min-Max]. See Fig. 4A and Fig. S4 to S8 for the demographic scenarios and parameters.

| **Priors for the demographic parameters** | |
|---|---|
| Effective size of the *i*-population ($N_i$) | UN~[100 – 10,000] |
| Splitting time ($T_j$, in generations) | UN~[10 – 1,000] |
| Duration of the bottleneck (*DB)* | UN~[1 - 10] |
| Effective size during the bottleneck ($NF_i$) | UN~[1 - 100] |
| Admixture rate $R_a$ | UN~[0 - 1] |
| Constraint on parameter | $T_k > T_j$ |

| **Priors for the mutation model for microsatellites** | |
|---|---|
| Mean mutation rate (MEAN - μ) | UN~[1 x $10^{-4}$ - 1 x $10^{-3}$] |
| Individual locus mutation rate (GAM - μ) | GA~[1 x $10^{-5}$, 1 x $10^{-2}$, 2] |
| Mean coefficient P (MEAN - P) | UN~[0.1, 0.3] |
| Individual locus coefficient (GAM - P) | GA~[1 x $10^{-2}$ , 9 x $10^{-1}$, 2] |
| Mean SNI rate (MEAN - SNI) | LU~[1 x $10^{-8}$, 1 x $10^{-5}$] |
| Individual locus SNI rate (GAM - SNI) | GA~[1 x $10^{-9}$, 1 x $10^{-4}$, 2] |
| Max. mutational steps | 40 |

**Table S14**. Summary statistics used in all DIYABC simulations.

| DIYABC abbreviation | Description |
|---|---|
| *Single sample statistics for each sampled population* | |
| NAL | Mean number of alleles across loci |
| HET | Mean gene diversity across loci (Nei, 1987) |
| VAR | Mean allele size variance across loci |
| MGW | Mean *M*-index across loci (Garza and Williamson, 2001; Excoffier et al., 2005) |
| *Two sample statistics for each pairwise sample combination* | |
| N2P | Mean number of alleles across loci (two samples) |
| H2P | Mean gene diversity across loci (two samples) |
| V2P | Mean allele size variance across loci (two samples) |
| FST | $F_{ST}$ between two samples (Weir and Cockerham, 1984) |
| LIK | Mean index of classification (two samples) (Rannala and Moutain, 1997; Pascual et al., 2007) |
| DAS | Shared allele distance between two samples (Chakraborty and Jin, 1993) |
| DM2 | $(\delta\mu)^2$ distance between two samples (Golstein et al., 1995) |
| *Admixture statistics for each combination of population 2 (WEUR), 3 (EEUR), 4 (CN) (only for step 1, Fig. S4)* | |
| AML | Maximum likelihood coefficient of admixture (Choisy et al., 2004) |
